# Prophage induction shifts community composition and functional capacity in a *Sargassum*-derived multispecies biofilm

**DOI:** 10.64898/2026.03.26.714470

**Authors:** Alexandra K. Stiffler, Natascha S. Varona, Bailey A. Wallace, Cynthia B. Silveira

**Affiliations:** Department of Biology, University of Miami, Coral Gables, FL 33146, USA; Department of Environmental Science, Policy, and Management, University of California, Berkley, Berkley, CA 94720, USA; Department of Marine Biology and Ecology, Rosenstiel School of Marine, Atmospheric, and Earth Science, University of Miami, Miami, FL 33149, USA

**Keywords:** Bacteriophage, Mitomycin C, microbiome, *Vibrio*, marine symbiosis

## Abstract

**Background:** Pelagic *Sargassum* has undergone significant range expansion and dramatic blooms in the Atlantic over the past 15 years. This algae’s microbiome provides symbiotic functions that are believed to contribute to its ecological success. Recent research shows that *Sargassum*-associated bacteria are enriched in integrated prophages compared to the surrounding seawater and that these prophages are inducible by chemical and ultraviolet treatment.

**Results:** Here, we investigated a *Sargassum*-derived *in vitro* multispecies biofilm encompassing the dominant heterotrophic microbial members associated with *Sargassum* to probe the impacts of prophage induction on the composition of *Sargassum* microbiomes. Induction was quantified by coverage-based virus-to-host ratios in chemically induced treatments with Mitomycin C and non-induced controls, and the community composition and metabolic profiles were analyzed after a period of recovery post-induction. Chemical induction led to a significant increase in abundance and virus-to-host ratio of viral genomes linked to *Vibrio* metagenome-assembled genomes. This was accompanied by altered biofilm community composition, with a reduction in *Vibrio* bacterial abundance that opened niche space for other biofilm members in the genera *Pseudoalteromonas*, *Alteromonas*, and *Cobetia*. The induced *Vibrio*-associated phages encoded genes involved in quorum sensing, biofilm formation, virulence, and host metabolism. Induction led to a relative loss of 17 metabolic modules, including functions related to energy metabolism and nitrogen utilization.

**Conclusion:** Due to the high frequency of lysogeny in the *Sargassum* microbiome and the susceptibility of prophages to chemical and ultraviolet light induction, these results suggest that prophage integration and induction are mechanisms that significantly contribute to structuring the *Sargassum* microbiome and its functional profiles, potentially aiding in microbiome flexibility in changing environmental contexts.

## Introduction

The macroalgae genus *Sargassum* comprises two species, *S. natans* and *S. fluitans*, which exhibit holopelagic life cycles, characterized by the absence of a benthic phase. Historically, pelagic *Sargassum*, herein referred to as *Sargassum*, was mainly restricted to the Sargasso Sea [1]. In this ultra-oligotrophic area of the North Atlantic, *Sargassum* rafts provide biodiverse hotspots for macrofauna [2–4] and fix and sequester carbon [5–7]. However, with the development of the Great Atlantic *Sargassum* Belt (GASB) across the Equatorial Atlantic since 2011, *Sargassum* biomass has increased drastically, reaching more than 20 million metric tons from the Gulf of Mexico to West Africa [8]. *Sargassum* super blooms have led to increased mass beaching events [9–11], with negative socioeconomic impacts on tourism and fishing, as well as potential health concerns and costly management [12–14]. These *Sargassum* blooms have also been linked to declines in seagrass beds and coral reefs due to reduced light and oxygen in nearshore water [15]. This unprecedented proliferation, along with the resulting ecosystem and socio-economic consequences, highlights the need to understand the mechanisms that have enabled *Sargassum*’s recent range expansion and growth.

Algal microbiomes include dense communities of epibiotic microorganisms that adhere to host surfaces by excreting extracellular polymeric substances, forming biofilms [16]. These biofilms have dynamic interactions with their algal host, often through nutritional symbiosis, in which both partners mutually benefit. Bacteria can utilize algal-derived oxygen, organic carbon [17, 18], and osmolytes such as dimethylsulfoniopropionate [19], putrescine, taurine [20], creatine, and sarcosine [21]. In return, the algal host receives vitamins and metabolites [22], such as B_12_ [23], fixed nitrogen [24], and antibacterial compounds to exclude opportunistic pathogens [25, 26]. In many cases, bacteria are essential for algal growth, as axenic algae in the genus *Ulva* do not develop properly, whereas the addition of bacterial strains can induce a morphology similar to that observed in natural growth [27, 28]. Alternatively, biofilms can negatively impact their algal hosts by favoring primary and opportunistic pathogens [29, 30]. Building on this existing knowledge of biofilms in other macroalgae, recent studies have begun to investigate the potential role of the *Sargassum* microbiome in supporting algal growth by providing novel nutrient sources, thereby aiding its ability to acclimate to a wide range of oceanic environments [31, 32]. While few studies are currently available, bacterial members of the *Sargassum* microbiome have been implicated in converting organic nitrogen [32, 33] and phosphorus [31] sources into bioavailable compounds, potentially supplying *Sargassum* with nutrient pools it would not otherwise be able to access. These studies indicate a central role of microbiomes in *Sargassum* physiology.

Across its range, *Sargassum*’s bacterial community is consistently dominated by the phyla *Pseudomonadota*, *Bacteroidota*, *Cyanobacteriota*, *Actinomycetota*, *Planctomycetota*, *Chloroflexota*, and *Verrucomicrobiota* [34]. Within *Pseudomonadota,* the most abundant phylum in the majority of *Sargassum* samples [34], the orders *Rhodobacterales*, *Alteromonadales,* and *Vibrionales* are frequently abundant [35–39]. However, microbiome composition varies with oceanographic context, with an increasing abundance of *Vibrionales* in the Caribbean and South Florida [38], in contrast to a dominance of *Alphaproteobacteria,* such as *Rhodobacterales,* in the Sargasso Sea and other regions where *Vibrionales* are less abundant [37]. These compositional differences are thought to reflect differences in microbiome function. *Rhodobacterales* are a metabolically diverse group with multiple trophic modes, contributing to vitamin production and to the synthesis and degradation of other metabolites [40]. *Vibrio* are heterotrophic bacteria with pathogenic strains capable of infecting humans and marine organisms; however, *Vibrio* species are found in a wide range of aquatic ecosystems and can be commensal and mutualistic members of microbiomes [41]. In *Sargassum*, *Vibrio* species were identified as the most abundant nitrogen-fixing bacteria [32] and have been shown to display pathogenicity-related phenotypes [42]. While these studies indicate an essential role of community composition in the *Sargassum* microbiome’s functions, the mechanisms regulating community structure are unknown.

In addition to bacteria, *Sargassum* biofilms host a diversity of bacteriophages (phages), viruses that infect prokaryotic organisms [35]. In seawater, phages can exert top-down control on bacteria by lysing their hosts and serve as a source of genetic diversity through horizontal gene transfer [43–45]. Temperate phages, which can integrate into their host’s genome as a prophage, encode genes that can impact their bacterial host’s growth, fitness, and virulence, in addition to protecting against superinfection by related phages [46–49]. Previous metagenomic analyses of *S. natans* in South Florida showed that the viral community of the algae is distinct from that of the surrounding seawater, and that *Sargassum*-associated bacteria harbor more prophages than seawater [35]. *Sargassum*-associated phages encode genes for biofilm formation, and laboratory experiments showed that prophage induction (excision from the host genome and subsequent cell lysis) decreased biofilm production, suggesting that prophages have the capacity to modulate the growth of *Sargassum* biofilms [35]. However, the impact of induction on the microbial community composition and function is unknown.

Here, we hypothesized that prophage induction can cause a significant change in community composition and function by lysing dominant community members, opening niche space for other members of the microbiome. We tested this hypothesis in a *Sargassum*-derived multispecies biofilm representative of natural *Sargassum* microbiomes, using *in vitro* chemical induction and shotgun metagenomics post-recovery.

## Methods

### Biofilm induction assay

The biofilm assay reported here was previously described in Stiffler *et al.* 2024, where it was initially conducted to test the inducibility of the integrated prophages, given that many prophages are domesticated and cannot be induced. Here, we quantify host and virus-specific induction and changes in community composition and function in these biofilms. Briefly, visually healthy *Sargassum* samples (*S. fluitans* and *S. natans*; Fig. 1a) were collected 2 m from the shore close to the University of Miami’s Rosenstiel School of Marine, Atmospheric, and Earth Science (RSMAES) campus on May 16th, 2023. About 150 g of *Sargassum* was collected from a single *Sargassum* patch in autoclaved containers, placed in a cooler, and transported back to the University of Miami’s main campus for immediate processing. The *Sargassum* was rinsed with DI water to remove loosely associated organisms, and the tissue was scraped with a cell scraper (Greiner Bio-One, Austria) to dislodge biofilms, which were collected into 200 mL of 100 kDa-filtered seawater. To further dislodge tightly associated microorganisms, algal tissue was placed in 250 mL tubes with 200 mL of 100 kDa-filtered seawater, vortexed for 20 minutes, and combined with the scraped biofilms. The biofilm suspension was filtered through a 10 µm filter mesh (Aquatic Experts, Greensboro, NC, USA) pre-treated with 10 % bleach and rinsed thoroughly with DI water. The filtered *Sargassum* biofilm suspension was centrifuged at 8000 x g for 30 minutes to pellet bacterial cells, and the supernatant, containing extracellular viruses and debris, was removed. Pellets were washed with 100 mL of 100 kDa-filtered seawater, re-pelleted, and resuspended in 100 kDa-filtered seawater containing 25% glycerol before storage at -80 °C for future use. We refer to this as the *Sargassum* bacterial enrichment. Cell counts of the *Sargassum* bacterial enrichment were determined using epifluorescence microscopy [35].

**Figure 1:**
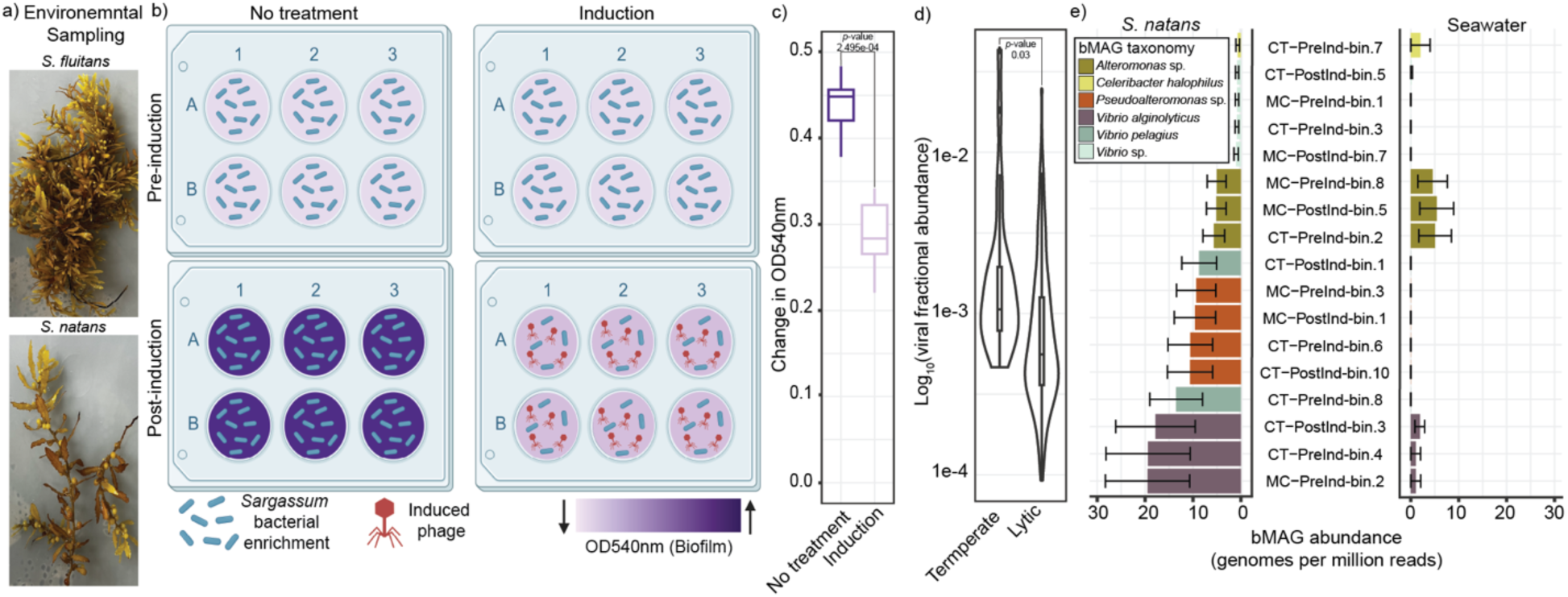
Mitomycin C treatment of a *Sargassum*-derived biofilm, representative of natural *Sargassum* heterotrophic bacterial members, decreases short-term biofilm formation capabilities. (a) Pictures of representative *S. fluitans* and *S. natans* samples collected for bacterial enrichment. (b) Cartoon representation of the *Sargassum*-derived biofilm induction experiment (Created in BioRender. Silveira, C. (2026) https://BioRender.com/m84ft3c). (c) Change in biofilm formation (quantified via OD540) of the *Sargassum* bacterial enrichment grown in 10% *Sargassum*-supplemented seawater 12 hours after induction for MC-treated and control samples. (Control *n* = 5, MC *n* = 5; ANOVA, *p* value = 1.69e-05; Tukey’s HSD, *p* value = 2.495e-04). (d) Abundance of temperate and lytic viruses recovered from biofilms in naturally occurring *Sargassum* (*n* = 213, shown as the log_10_ median fractional abundance across five replicates). (e) Abundance of the 17 bMAGs from biofilms that had greater than 0.5 genomes per million reads in in situ *Sargassum* samples (mean, *n* = 5) compared to their relative abundance in the surrounding seawater.

To determine the impact of prophage induction on *Sargassum*-derived biofilms, the *Sargassum* bacterial enrichment was grown in a 10 % *Sargassum*-supplemented seawater solution. To prepare the *Sargassum*-supplemented seawater, 100 g of mixed samples *S. fluitans* and *S. natans* were ground with a mortar and pestle, blended with 1 L of 0.45 µm filtered seawater, filtered through a 100 µm mesh and a 0.45 µm Sterivex filter (MilliporeSigma, Burlington, MA, USA), and autoclaved. For the biofilm assay, 5 mL of 10 % *Sargassum*-supplemented seawater was inoculated with the *Sargassum* bacterial enrichment at a final concentration of 1.4 × 10^6^ cells/mL in eight individual 6-well plates (CytoOne, USA Scientific), with one well in each plate used as a blank. Four plates were used for optical density (OD) quantification at 540_nm_, and the remaining four were harvested for sequencing. The plates were divided into two groups: a control group with no treatment and an induction group that would be treated with 1 µg/mL of Mitomycin C (herein referred to as MC; Fig. 1b). All plates were incubated at room temperature (25 °C) for 12 hours.

After 12 hours, two replicate plates (each with five independent replicate wells and one technical control well) from each group were subject to OD_540_ quantification or biofilm harvesting (pre-induction time point). For OD_540_ quantification, the media was removed, and the wells were rinsed twice with 1 mL of 0.02 µm filtered artificial seawater. The biofilms were stained with 2 mL of 0.01% crystal violet (Eastman Organic Chemical, Rochester, NY, USA) for 10 minutes and subsequently washed three times with 2 mL of 0.02 µm filtered artificial seawater. For quantification, 2 mL of 100% ethanol was added to each well, and OD_540_ was measured using a Synergy H1MD Hybrid Reader (BioTek, USA).

For sequencing, biofilms were harvested by removing the supernatant, adding 500 µL of 0.02 µm filtered artificial seawater, and scraping the biofilms off with a cell scraper. Samples were stored at -80 °C until DNA extraction. At this time, the 10% *Sargassum*-supplemented seawater was changed for the remaining plates, MC was added to the induction group, and plates were left to grow for an additional 12-hour recovery period before repeating OD_540_ quantification and harvesting biofilms.

### DNA Extractions and Sequencing

DNA extractions of the biofilm samples were performed with a DNeasy Blood and Tissue Kit (QIAGEN, Germantown, USA). DNA concentrations were quantified using a Qubit 2.0 Fluorometer using a High-Sensitivity dsDNA kit (Thermo Fisher Scientific, Waltham, USA).

DNA library preparation and sequencing were performed by AZENTA, Inc. in South Plainfield, NJ, USA, using the NEBNext Ultra DNA Library Preparation kit according to the manufacturer’s instructions (NEB, Ipswich, MA, USA). The genomic DNA underwent fragmentation using acoustic shearing via a Covaris S220 and was subjected to end-repair and subsequent adapter ligation following the adenylation of 3′ ends. Adapter-ligated DNA was indexed and enriched by limited-cycle PCR. The resulting libraries were verified using TapeStation (Agilent Technologies, Palo Alto, CA, USA) and quantified using Qubit 2.0 Fluorometer and real-time PCR (Applied Biosystems, Carlsbad, CA, USA). Sequencing was performed on an Illumina HiSeq 4000 (Illumina, San Diego, CA, USA) in a 2 × 150 paired-end configuration. The base calling was performed using the HiSeq 4000 or equivalent Control Software, and the bcl files were converted to fastq and demultiplexed using bcl2fastq v2.17.

### Quality control and co-assemblies of metagenomic data

Shotgun sequencing yielded an average of 3.2 × 10^7^ raw reads per sample (Table S1), showing the number of reads before and after quality control. The reads were filtered and trimmed using BBduk v39.06 [50] with the following parameters: left and right trimming (qtrim=rl, trimq=30), adapter trimming of both ends with a k-mer size of 23 (ktrim =rl, k=23, mink=11), a hamming distance of one, and tpe/tbo parameters. Read quality was quantified with FastQC v0.11.9 [51], and reads passing QC were co-assembled based on treatment (*n* = 5 per treatment) using MEGAHIT v1.2.9 [52], which generated 61,583, 48,143, 20,026, and 28,699 contigs for control pre-induction, MC pre-induction, control post-induction, and MC post-induction, respectively.

### Viral genome identification and curation

Viral sequences were identified among the co-assembled contigs using geNomad v1.8.0 [53]. GeNomad employs a neural network model to identify nucleotide patterns, utilizing an encoder that contains a framework based on IGLOO architecture [54]. It then predicts open reading frames using a modified version of Prodigal [55] and queries predicted proteins against a custom set of plasmid and virus-specific protein profiles using MMseqs2 [56]. Viral contigs identified using geNomad were extended using COBRA v1.2.3 [57]. Briefly, COBRA assesses breakpoints in the assembly de Bruijn graph and uses coverage and paired-read spanning to determine whether contigs should be extended. Reads were mapped at 95% identity to co-assembled contigs within each treatment group using SMALT v0.7.6 [58] for COBRA analysis. SAM (Sequence Alignment/Map) files were sorted and indexed using SAMtools v1.10 [59].

Coverage depth files were generated from the mapping files using the Joint Genome Institute’s jgi_summarize_bam_contig_depths script, and the coverage file was formatted for COBRA using their coverage.transfer.py script. COBRA was run with these input files and maxk and mink parameters. Extended contigs were analyzed through geNomad for provirus identification and updated viral taxonomic classification. GeNomad viruses were dereplicated using Virathon (GitHub - Felipehcoutinho/Virathon: Genomic Analysis of Viruses of Archaea and Bacteria). Briefly, Virathon creates viral populations using Prodigal v2.6.3 [55] to identify open reading frames, count genes per sequence, and perform an all-vs-all BLASTn [60] comparison at the gene level to calculate the percentage of shared genes and ANI. Viral genomes with 80% shared genes and 95% ANI are considered to be within the same population, and a greedy approach is used to retain the longest genome as the population representative. The quality of the dereplicated viral sequences was then assessed using CheckV v1.0.1 [61]. Viral sequences classified as medium/high quality, longer than 10 Kb, or as a prophage with a flanking region greater than 2000 bp by geNomad or CheckV were retained, yielding 334 viral genomic sequences for further analysis. The infection strategy, temperate or lytic, for these 334 viruses was predicted using DeepPL [62], which utilizes DNABERT [63], a pre-trained bidirectional encoder representation trained on a dataset with equal representation of nucleotide sequences encoding temperate and lytic genes. Viral sequences were annotated using MetaCerberus v1.3.1[64], which employed Prodigal and all available HMMs.

### Identifying prophage induction using coverage-based virus-to-host-ratio

To determine if the identified prophages were induced in MC-treated samples, coverage data of prophage regions was compared to that of bacterial flanking sequences to generate a coverage-based virus-to-host-ratio (VHR) for each prophage in each sample using PropagAtE v1.1.0 [65]. First, reads were mapped at 99% identity to the original contigs, in which the prophages were identified at 99% identity using SMALT v0.7.6 [58]. SAM (Sequence Alignment/Map) files were sorted using SAMtools v1.10 [59]. QC paired read files, contigs with prophage and host regions, SAM, sorted BAM, and a prophage coordinates file (obtained from geNomad and CheckV) were formatted for PropagAtE v1.1.0 input. We note that PropagAtE commonly uses a VHR of 1.5 as a threshold for identifying induction. However, in complex metagenomic samples, not all bacterial hosts are lysogenized, and not all prophages will simultaneously induce, which can potentially lower this ratio (here, 63% of VHRs were below 1). Here, we leveraged our study design, in which bacterial biofilms were harvested before and after induction with a known inducing agent, to calculate VHR pre- and post-induction and identify induced prophages as those with a statistically significant increase in VHR, even when values were below 1.5.

### Identifying putative induction of viruses using viral fractional abundances

In addition to the induction of prophages identified with clear bacterial flanking regions, we investigated the putative induction of phages that could not be assembled with bacterial flanking regions, which could be caused by multiple integration sites [66, 67] or a small fraction of the host population being lysogenized [68]. The fractional abundance of all identified viral sequences was calculated by mapping the reads from each sample to the biofilm viral database at 95% identity using SMALT v0.7.6. SAM files were sorted and indexed with SAMtools v1.10.

Fractional abundances (𝑓_(*i*)_) were calculated by taking into account contig length using the equation below [69].

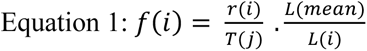

where (r_(*i*)_) is the number of reads mapped to each viral genome sequence obtained with SAMtools v1.10, L(*mean*) corresponds to the mean viral genome length (bp) in the database, L(*i*) denotes the length of the viral genome, and (T(*j*)) represents the number of viral mapped reads in each metagenome sample.

### Binning and analysis of bacterial metagenome-assembled genomes

Bacterial metagenome-assembled genomes (bMAGs) were constructed by mapping QC reads to co-assembled contigs using Bowtie2 v2.3.5 [70]. Using SAMtools v1.10 [59], the SAM files were compressed, sorted, and indexed according to the binning algorithm specifications. Three binning programs were employed to build bMAGs: CONCOCT v1.0 [71], MetaBAT2 v2.12.1 [72], and MaxBin2 v2.2.6 [73]. All three binners combine contig coverage and sequence composition to cluster contigs into genomes. The following raw bins were refined, reassembled, and quantified using the *bin_refinement*, *reassemble_bin,* and *quant_bins* modules of MetaWRAP v1.3.2 [74]. One replicate sample from the control post-induction group contained a bMAG that matched *Salmonella*, a common contaminant from human material. As this bMAG was assembled with sequences coming from a single replicate, and NMDS analysis indicated it was an outlier (Fig. S1), this bMAG was treated as a contaminant and removed from the study. The remaining bMAG relative abundances in the replicate sample where this bMAG originated were not distinguishable from the other replicates, so this metagenome was maintained for subsequent analyses after the removal of this bMAG (Fig. S1). Bin abundance was obtained through MetaWRAP in genomes per million reads using a modified version of SALMON v0.13.1 [75]. This abundance estimate considers a length-weighted bMAG coverage of individual contigs relative to the number of reads per sample. CheckM2 v1.0.2 [76] estimated bMAG completion and contamination. Taxonomy was assigned to bMAGs using GTDB-tk v2.1.1 [77, 78]. KEGG metabolism pathways were assigned to bMAGs using the *anvi-run-kegg-kofams* and *anvi-estimate-metabolism* modules of Anvi’o v8 with default settings [79]. Only modules that had a pathwise completeness of 70% or greater were retained for further metabolism analysis. bMAGs were clustered based on average nucleotide identity (ANI) using *anvi-dereplicate-genomes* module [79] with fastANI v1.32 using the ‘*min-full-percent-identity*’ and ‘*similarity-threshold*’ flags at 97% ANI [80]. The Newick tree file was visualized and annotated using iTOL v7 [81].

### Host prediction

Prophages were linked to bMAGs by identifying contigs containing prophages within the bin. One bMAG (CT-PostInd-bin.5) had a viral sequence (CT-PostInd-BF1_k127_431539) binned within it, but with no flanking regions. This viral sequence was part of the same viral population as an identified prophage (MC-PostInd-BF1_k127_567814; prophage.8). We assume that the prophage identified within the MC post-induction samples would not have been binned with its host due to induction-related coverage shifts. Therefore, BLASTn [60] was used to align the bacterial flanking region of the prophage to the non-prophage viral sequence within bMAG CT-PostInd-bin.5. The last 648 bp of the viral sequence matched the first 648 bp of the bacterial flank at 99.3% identity; thus, we deemed this a prophage-bMAG link (Table S2).

Bacterial flanks surrounding prophages (>2000 bp) were extracted and compared to the GTDB GCF representative genomes (accessed September 2025) [78] using BLASTn [60]. Only matches that had greater than 85% identity and an e-value less than 1e-10 were considered (Data S5). For a genus to be assigned to a bacterial flank 50% of the blast hits had to be assigned to a single genus. Additionally, to predict the genus of the host for viruses in the biofilm induction database, we used iPHoP v1.3.3 [82] using the ‘Aug_2023_pub’ host database and default ‘predict’ parameters. iPHoP combines six different methods to predict phage-host pairs. Using neural networks and random forest classifiers, it computes hit scores for potential hosts by combining the host-based prediction tools BLAST [60], CRISPR, VirHostMatcher [83], WIsH [84], and PHP [85]. It then couples this with the phage-based prediction tool RaFAH [86] to assign genus or genome-resolved phage-host pairs. If methods predicted different hosts, both matches were discarded from the results but still recorded in supplemental datasheets (Table S3 & Table S4).

### Analysis of prophage and host-encoded AphA transcriptional regulator

One prophage (prophage.8) that was linked to a bMAG encoded the *aphA* gene, which produces AphA, a protein involved in biofilm formation and quorum sensing in several *Vibrio* species [87]. Interestingly, *aphA* was also identified as a potential biofilm gene encoded by a phage within the *S. natans* microbiome [35]. Therefore, we investigated the similarity of these phage and host-encoded proteins at the sequence and structural levels. MetaCerberus v1.3.1 [64] was used to annotate the bMAG host of the prophage.8 (CT-PostInd-bin.5; *Vibrio* sp.) and viral sequences identified in *S. natans* metagenomes from our previous study [35]. CT-PostInd-bin.5 and one viral sequence (sarg_contig.k127_668789) encoded genes for the AphA protein. A BLASTp [60] against the RefSeq non-redundant protein database [88] was used to identify homologous proteins to the *Vibrio* host and prophage.8 proteins. The 25 matches with the highest identity and coverage were selected for each, dereplicated at 99% identity using CD-HIT v4.8.1 [89], and compared with the two AphA amino acid sequences from this study and the one viral sequence from the *S. natans* metagenomes, for a total of 40 amino acid sequences, via alignment with MAFFT v7.511 [90]. A phylogenetic tree of amino acid sequences was constructed using the Neighbor-Joining algorithm with the conserved sites method and the JTT substitution model with 100 bootstrap resampling [90]. The Newick tree file was visualized with iTOL v7 [81]. The tridimensional structure of the AphA proteins encoded by prophage.8 and CT-PostInd-bin.5 were predicted with AlphaFold 3 [91]. Predicted structures were aligned using the *matchmaker* command of ChimeraX v.1.10.1 [92].

### In situ abundances of biofilm induction viruses and bMAGs

The presence of biofilm induction viruses and bMAGs in *S. natans* and seawater samples collected off the coast of Florida was determined by mapping QC reads from *S. natans* and seawater to the viral sequences identified in this study at 95% identity using SMALT v0.7.6 [58]. SAM files were sorted and indexed using SAMtools v1.10 [59], and coverage was obtained using BBMap’s ‘pileup.sh’ script [50]. For a virus to be labeled as present in the *S. natans* and seawater samples, it had to have greater than 75% coverage. Viral fractional abundance was calculated using the previously described equation. Relative abundance of the 28 biofilm induction bMAGs in the same *S. natans* and seawater samples was quantified using the *quant_bins* module of MetaWRAP v1.3.2 [74]. bMAGs with fewer than 0.5 genomes per million reads in *S. natans* samples were considered absent.

### Figures and statistics

All figures and statistical analyses were generated using R v4.3.2, and aesthetic edits were made in Adobe Illustrator 2025. Data spreadsheets were read using readxl v1.4.5 [93], and data sheets were re-formatted when needed with tidyr v1.3.1[94], dplyr v2.5.0 [95], tidyverse v2.0.0 [96], and stringr v1.5.1 [97]. The non-metric multidimensional scaling (NMDS), permutational multivariate analysis of variance (PERMANOVA), and similarity percentage (SIMPER) analyses were conducted using vegan v.2.7-1 [98]. NMDS analyses were computed using Bray-Curtis dissimilarity matrices with 9999 permutations, and pairwise PERMANOVA analyses were performed using pairwiseAdonis v0.4.1 [99]. Bar plots, boxplots, pie charts, and NMDS plots were generated with ggplot2 v3.4.1 [100], ggExtra v0.10.0 [101], and ggpubr v0.6.0 [102]. Heatmaps were created with pheatmap v1.0.12 [103], viral genome maps with genoPlotR v0.8.1 [104], and bMAG percent nucleotide identity, and the protein phylogeny trees were visualized and edited with iTOL v7 [81].

## Results

Mitomycin C (MC) treatment of the *Sargassum*-derived biofilm communities reduced biofilm growth, as measured by the change in OD_540_ between pre- and post-induction, where a significant decrease in the MC-treated group was observed compared to the untreated control (ANOVA, *p* value = 1.69e-05; Tukey’s HSD, *p* value = 2.495e-04; Fig. 1c). An initial analysis of biofilm viral communities in this experiment showed that 33% of the putative phages identified increased in fractional abundance in the MC group. Among the phages with increased abundance, 19.25% were categorized as temperate in the MC-treated group, compared to only 3.71% in the control. These results indicated the presence of induced prophages that went undetected by phage identification tools used in a previous study [35]. By applying viral contig extension [57], an additional identification tool [53], and screening bacterial metagenome-assembled genomes (bMAGs), we identified several putatively induced phage-host pairs, as described below.

Binning of contigs resulted in 28 bacterial metagenome-assembled genomes (bMAGs) that were > 50% complete (mean = 80.49%, SD = 14.82%) and < 10% contaminated (mean = 2.39%, SD = 2.12%; Data S1). We identified 1,303 putative viral genomes/genome fragments that were categorized as medium or high quality, longer than 10 Kb, or labeled as a prophage with a bacterial flanking region longer than 2,000 bp. These were dereplicated into 334 viral populations (herein referred to as viruses; Data S2).

### Sargassum-derived biofilms are representative of naturally occurring Sargassum microbiomes

To assess environmental relevance, *S. natans* and seawater metagenomic reads collected off the coast of South Florida [35] were mapped to the viruses and bMAGs from this study. Of the 334 viruses derived from the biofilm assay, 64 % were present in *S. natans* metagenomes (Fig. S2), but none were identified in surrounding seawater metagenomic reads. These overlapping viruses consisted of two prophages, 25 temperate phages, and 186 putatively lytic viruses (Fig. S2). The majority of these 213 viruses were unclassified (79.81%), with the remaining viruses belonging to *Pokkesviricetes* (10.33%), *Revtraviricetes* (5.63%), *Caudoviricetes* (3.29%), and *Faserviricetes* (0.94%) (Fig. S2). The 27 temperate viruses (including the two prophages) were more abundant in *S. natans* compared to the lytic viruses (Fig. 1d; Wilcoxon rank sum: *p*-value = 0.03). The three most abundant biofilm viruses in *S. natans* environmental samples included two temperate phages (MC-PostInd-k127_604294 and prophage.8) and one lytic virus (CT-PreInd-k127_631518) (Data S3). Within environmental *S. natans* samples, these three viruses accounted for 7.5%, 3.2%, and 4.2% of the biofilm assay viral community identified, respectively.

All bMAGs from the biofilm induction experiment together had, on average, 138.86 genomes per million reads in *S. natans* samples (Fig. 1e). An analysis of the relative abundance of these bMAGs in the *S. natans* samples demonstrated that 17 of the 28 bMAGs were present in environmental *Sargassum* samples. The most abundant bMAG, classified as *Vibrio alginolyticus*, had an average of 19.46 genomes per million reads, accounting for 10% of the total number of genomes (Fig. 1e; Data S4). Only nine of the 28 bMAGs were present in seawater, indicating that the *in vitro* biofilm was less representative of the seawater microbial community, as expected. Seven bMAGs were found in both *S. natans* and seawater samples: one *Celeribacter halophilus*, three *Alteromonas* sp., and three *Vibrio alginolyticus*. The ten bMAGs unique to *S. natans* included two *Vibrio pelagius*, four *Vibrio* sp., and four *Pseudoalteromonas* sp.

### Mitomycin C induces Vibrio prophages and affects viral abundance

Of the 334 viruses, 11 had representatives identified as prophages with an average length of 40,502 bp and an average flanking region length of 12,550 bp (Table S3). Eight prophages belonged to the class *Caudoviricetes*, one in the family *Autographiviridae*, one in *Kyanoviridae*, and one in *Corticoviridae* (Fig. 2a). Seven prophages were predicted to infect the genus *Vibrio* (Table S3). Virus-to-host ratios (VHRs) (Data S6) were similar between pre- and post-induction controls (Pairwise PERMANOVA: *p*-value = 0.62), as expected, and differed between MC pre-induction samples compared to post-induction samples (Fig. 2b; Pairwise PERMANOVA: *p*-value = 0.015). SIMPER analysis identified four prophages that accounted for most of the difference between the MC pre- and post-induction samples (Fig. 2b). Two of these prophages also differed between the control and MC post-induction samples, even though a comparison accounting for all prophages was not significant, indicating heterogeneity in prophage response to chemical induction (Pairwise PERMANOVA: *p*-value = 0.35). The four prophages identified in the MC post-induction were taxonomically classified as *Caudoviricetes* (two without family-level classification), one as *Autographiviridae*, and one as *Corticoviridae*. All were predicted to infect bacteria in the genus *Vibrio* (Fig. 2c). Two prophages, whose VHRs drove differences in both comparisons, encoded genes for host evasion, viral defense, and a transcriptional regulator belonging to the PadR family. Prophage.8 encoded *hdf* and *ulx*-like genes, which are a part of antirestriction systems to evade host defenses [105] and a transcriptional regulator with high sequence similarity to *aphA*, a regulator of quorum sensing and biofilm formation in multiple *Vibrio* species [106, 107] (Fig. 2c; Data S7). Furthermore, Prophage.10 encoded a gene with similarity to *terB*, which is involved in tellurite resistance and also plays roles in host antiviral defense systems [108].

**Figure 2:**
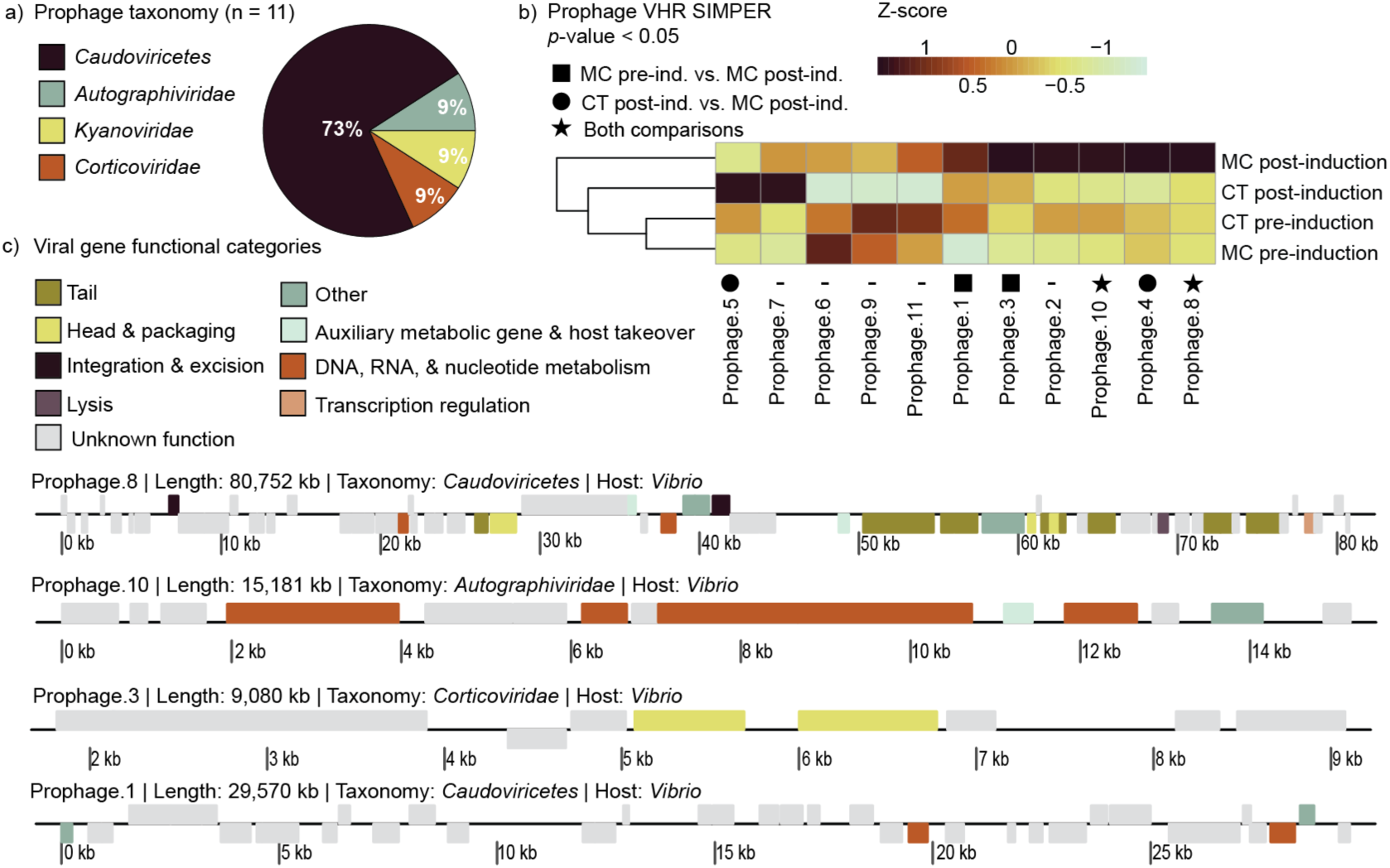
Mitomycin C induces pelagic *Sargassum* biofilm-associated prophages. | (a) Taxonomy of identified prophages (*n* =11); *Caudovircetes* (dark purple), *Autographiviridae* (blue), *Kyanoviridae* (light green), and *Corticoviridae* (orange). (b) Virus-to-host ratio (VHR) for each prophage, normalized by prophage with a Z-score (median, *n* = 5 for each sample types). Prophages marked with a square (MC pre- vs. MC post-induction), circle (CT post- vs. MC post-induction), or star (both comparisons) were identified by SIMPER analysis as significantly contributing to the differences between samples (*p* < 0.05). (c) Genome plots of the four prophages contributing to differences in the MC pre- and post-induction samples. Annotations for the plots are available in Data S7.

Most viral sequences in this study could not be taxonomically classified (61.4%). The remaining viruses belong to the following classes: *Caudoviricetes* (25.1%), *Pokkesviricetes* (7.2%), *Revtraviricetes* (4.5%), *Tectiliviricetes; Corticoviridae* (0.9%), *Faserviricetes; Inoviridae* (0.6%), and *Megaviricetes* (0.3%) (Fig. 3a). The abundances of these viruses (Data S8) differed significantly between MC and control pre- and post-induction (Pairwise PERMANOVA: *p*-values = 0.0067 and 0.0096 for MC pre vs. post induction and control pre- vs. post-induction, respectively) and post-induction groups (Pairwise PERMANOVA: *p*-value = 0.0066 control post vs MC post-induction). Pre-induction groups did not differ (Pairwise PERMANOVA: *p*-value = 0.15 control pre- vs. MC pre-induction) (Fig. 3b). SIMPER analysis identified 110 viruses driving the differences in viral abundance between MC and control samples. The change in fractional abundance in MC and control samples was calculated for these viruses to identify potential viral induction among viruses without bacterial flanking regions, which are incontrovertible evidence of integrated prophages, but are difficult to assemble when prophages integrate in different regions of the genome, or when a phage remains as an episome without genome integration. Twenty-one viruses displayed a positive shift in abundance that was greater in MC samples than in controls (Fig. 3c). Eight of these induction candidates were categorized as temperate phages encoding lysogeny-related genes (Fig. 3c): integrase, antitermination protein Ant, regulatory protein Rha, recombination protein Bet, recombination protein RecE, Cl repressor, antitermination protein Q, and replication protein O (Fig. 3d).

**Figure 3:**
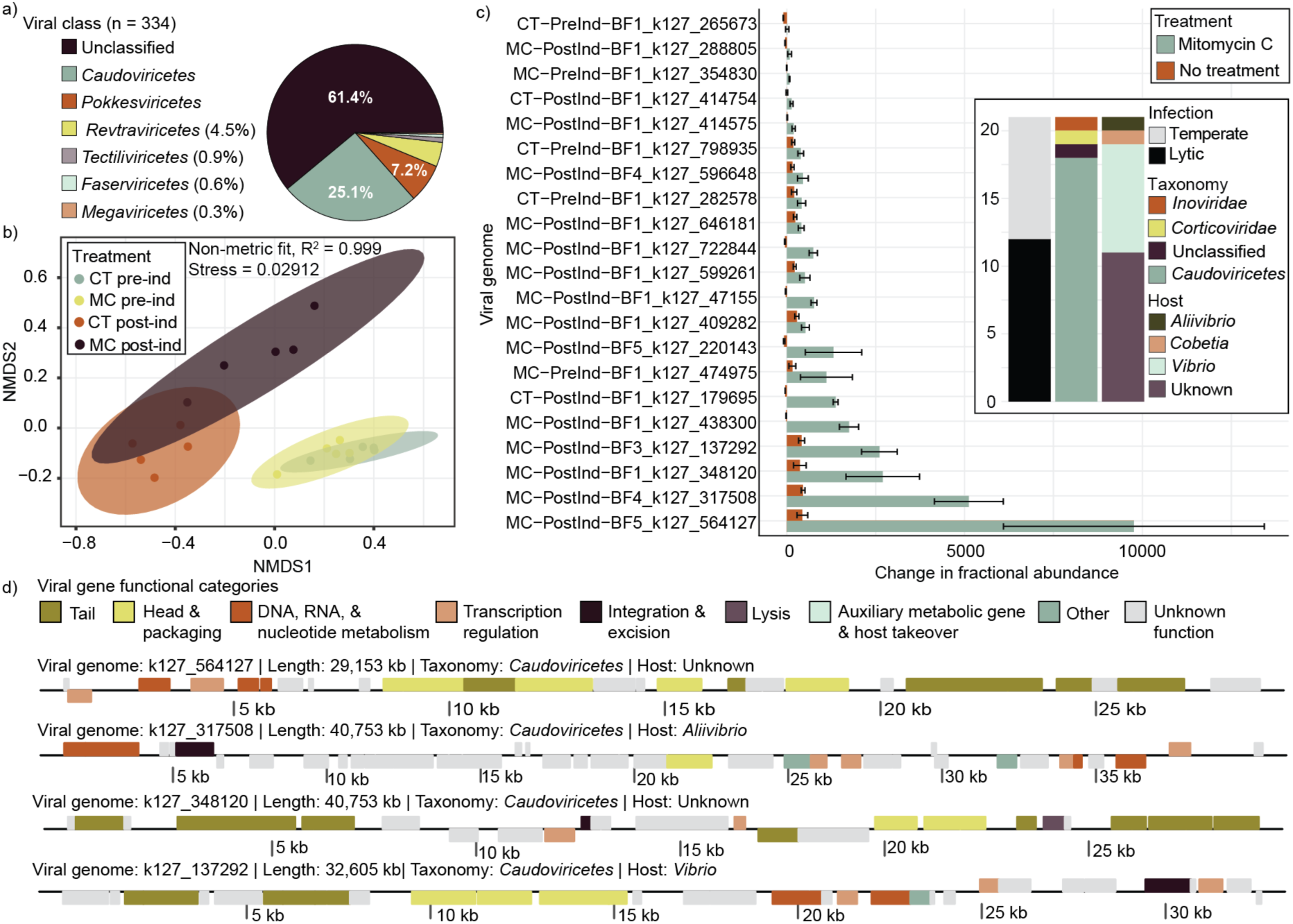
Viral fractional abundance changes upon induction, and temperate phages display a significant increase. (a) Taxonomy of all viruses identified in the biofilm (n = 334 after quality thresholds described in the methods); unclassified (dark purple), *Caudovircetes* (blue), *Pokkesviricetes* (orange), *Revtraviricetes* (light green), *Tectiliviricetes* (light purple), *Faserviricetes* (light blue), and *Megaviricetes* (light orange). (b) Non-metric Multidimensional Scaling (NMDS) of the viral fractional abundance across sample types, CT post-induction (*n* = 5, orange), CT pre-induction (*n* = 5, light blue), MC post-induction (*n* = 5, dark purple), and MC pre-induction (*n* = 5, light green). Each dot represents a replicate. The ellipses indicate the 95% confidence interval around each sample type. (c) Median change in viral fractional abundance for the 21 non-prophage viral induction candidates (SIMPER: *p* < 0.05). The colors represent the different treatments: no treatment (orange) and MC (blue). The error bars indicate standard error. (d) Genome plots for the four viruses that had the most significant change in fractional abundance between treatment groups. Annotations for the plots are available in Data S7.

Eighteen of these viruses were classified as *Caudoviricetes*, one as *Corticoviridae,* one as *Inoviridae*, and the remaining virus was unclassified. Ten of the 21 induction candidates had a predicted host. Eight viruses were predicted to infect *Vibrio*, one *Aliivibrio*, and one *Cobetia*. The *Inoviridae* phage was predicted to infect *Vibrio* and encoded zonula occludens toxin (Zot) (Data S7). Viral genome k127_317508, which is a temperate phage predicted to infect *Aliivibrio* and experienced the second greatest change in abundance between groups, encoded a chitinase (Fig. 3d; Data S7).

### Biofilm community composition shifts in response to induction

To investigate the impact of prophage induction on biofilm composition after 12 hours of recovery, we analyzed the abundances and genomic profiles of the 28 bMAGs identified in our dataset (Figure 4). The bMAGs comprised 17 *Vibrio*, four *Pseudoalteromonas*, three *Cobetia*, three *Alteromonas*, and one *Celeribacter*. All of these are a part of common bacterial families in *Sargassum* (*Vibrionaceae*, *Halomonadaceae*, *Alteromonadaceae*, and *Rhodobacteraceae*) [35–37]. Six bMAGs could be classified to the species level, with three *Vibrio alginolyticus*, two *Vibrio pelagius*, and one *Celeribacter halophilus* (Data S1). The relative abundance of the bMAGs, in genomes per million reads, differed significantly between sample types (Fig. 4a; PERMANOVA: *p*-value = 1e-04). The two pre-induction groups did not differ (Pairwise PERMANOVA: *p*-value = 0.22), the post-induction groups varied from their pre-induction counterparts (Pairwise PERMANOVA: *p*-value = 0.0085 and 0.0078 for control and MC, respectively), and the variation between the two post-induction groups was significant (Pairwise PERMANOVA: *p*-value = 0.025). SIMPER analysis identified 18 bMAGs that were differentially abundant when comparing the MC samples pre- and post-induction, and between control and MC post-induction (Fig. S3 depicts SIMPER *p*-values for all comparisons of the 18 significant bMAGs; Data S1). Among the 18 bMAGs driving differences between treatment groups, 13 *Vibrio* bMAGs decreased in abundance in the induced samples (Fig. 4b).

**Figure 4:**
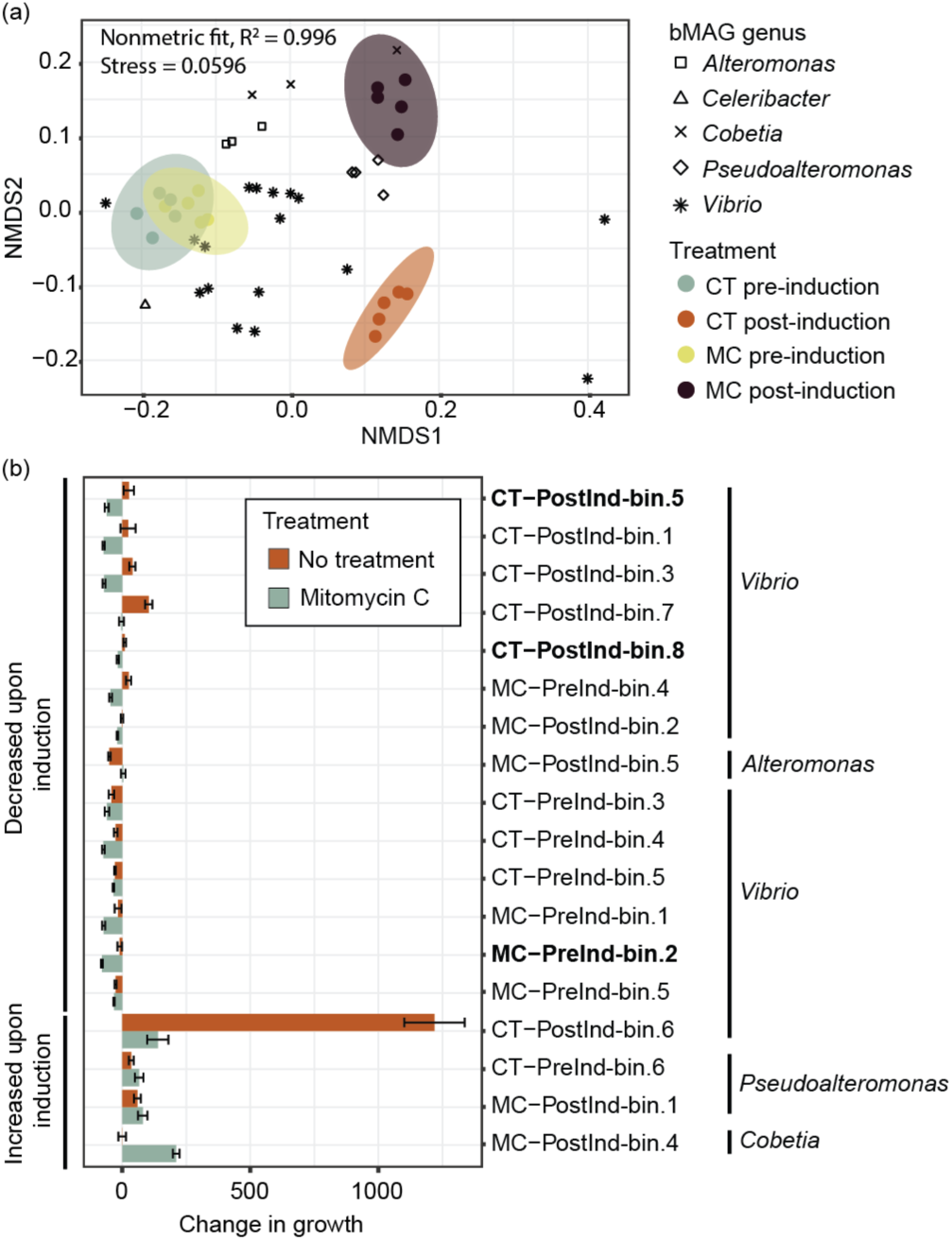
Bacterial community composition shifts upon prophage induction. (a) NMDS analysis of bMAGs across CT post-induction (*n* = 5, orange), CT pre-induction (*n* = 5, light blue), MC post-induction (*n* = 5, dark purple), and MC pre-induction (*n* = 5, light green). Each dot represents a replicate. The genus of each bMAG is defined by the shapes: *Alteromonas* (square), *Celeribacter* (triangle), *Cobetia* (x), *Pseudoalteromonas* (diamond), and *Vibrio* (star). The ellipses indicate the 95% confidence interval around each sample type. (b) Median change in abundance for the 18 bMAGs driving differences between the treatments (SIMPER: *p* < 0.05). The colors represent the different treatments: no treatment (orange) and MC (blue), error bars indicate standard error, and bolded bMAGs indicate a prophage link.

*Pseudoalteromonas* and *Cobetia* bMAGs became more abundant in the induced samples. One *Alteromonas* bMAG did not experience a change in growth in MC samples, but decreased significantly in the control (Fig. 4b and Fig. S3). Among the bMAGs decreasing in abundance with induction, two *Vibrio* sp. bMAGs (CT-PostInd-bin.5 and CT-PostInd-bin.8) and one *Vibrio alginolyticus* (MC-PreInd-bin.2) encoded prophage.8 prophage.3, and prophage.9, respectively. Prophage.8 encodes a transcriptional regulator with similarity to a gene that corresponds to the protein AphA and is a homolog of a host-encoded version of the protein. AphA belongs to the PadR superfamily, which is characterized by a winged helix-turn-helix motif. In many *Vibrio* species, it has central roles in quorum sensing, biofilm formation, and virulence[109]. CT-PostInd-bin.5’s AphA amino acid sequence was most like that of *Vibrio scophthalmi* (100% identity with 96% coverage; Data S9). The prophage amino acid sequence was most similar to a *Vibrio* multispecies version of AphA, meaning a non-redundant version of the protein found in multiple different species (99.37% identity with 99% coverage; Fig. 5a; Data S9). The AphA proteins of prophage.8 and its host only shared 34.78% identity (Table S5) at the amino acid level but displayed high structural similarity with a similarity score of 302.3 and RMSD between 82 pruned atoms of 1.216 Å (Fig. 5b; Fig. S4). This prophage-encoded protein was also observed in one viral sequence identified in the *in situ* microbiome of *S. natans* (sarg_contig.k127_668789), with 99.34% identity and 100% coverage (Fig. 5a; Table S5). Both phage-encoded *aphA* (phrog_13512) genes are flanked by viral genes (Ref-like RecA filament endonuclease, uncharacterized viral protein, and a probable phage tail protein) (Fig. S5; Data S7).

**Figure 5:**
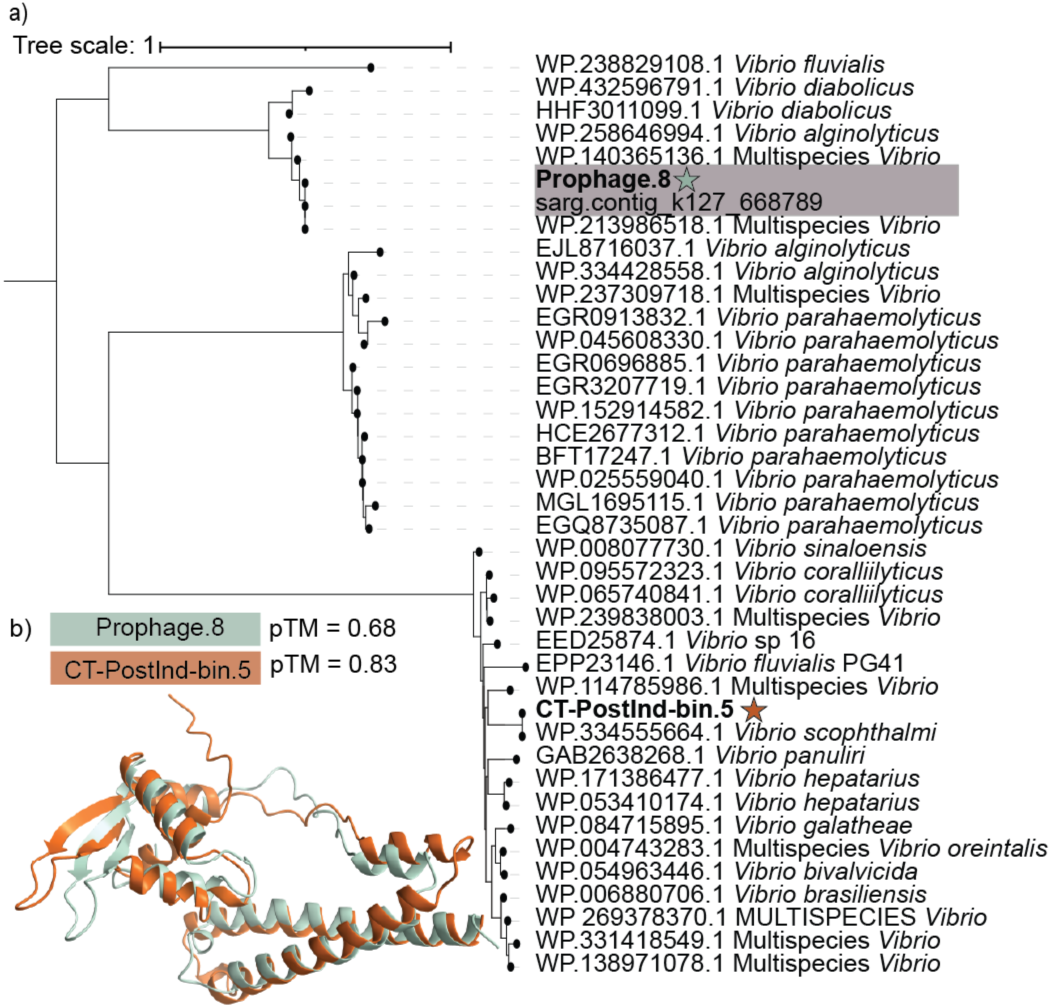
Prophage- and host-encoded AphA are divergent but have structural similarity. a) A Neighbor-Joining tree of 40 AphA amino acid sequences, two from this study (indicated by stars), one encoded by a viral sequence identified from *S. natans* metagenomes, and the top 25 RefSeq matches to both proteins dereplicated at 99% identity (n= 37). The light purple box indicated proteins encoded by viral sequences. b) Aligned AphA protein structures for prophage.8 (blue) and CT-PostInd-bin.5 (orange). Protein structures are denoted by a star on the phylogenetic tree with the colors above. The sequence alignment score between the two proteins was 302.3, and the RMSD between 82 pruned atom pairs was 1.216 Å.

### Prophage induction alters bacterial biofilm metabolic potential

Bacterial MAGs encoded 132 KEGG modules with a pathwise module completion of greater than 70 % in 26 submodules (Fig. 6a; Data S10). Ten submodules were present in all bMAGs (Fig. 6a). To investigate if induction shifted the microbiome metabolisms, the relative abundance of each bMAG that encoded a metabolism module was summed within each replicate as an indicator of module abundance. SIMPER analysis identified 17 modules that differed between the MC and control post-induction groups (Fig. 6b) and 108 modules between the MC pre- and post-induction. Here, we focus on the differences between the MC and control post-induction samples, as they provide insight into which metabolic pathways were depleted by prophage induction; no modules were enriched. The 17 modules were categorized into 10 different submodules: ATP synthesis, nitrogen metabolism, cofactor and vitamin metabolism, metabolic capacity, serine and threonine metabolism, polyamide metabolism, other aromatic metabolism, histidine metabolism, lipid metabolism, and other amino acid metabolism. The modules that decreased in MC treated compared to the control post induction groups were GABA biosynthesis, acetogen, ectoine degradation, cobalamin biosynthesis, thiamine biosynthesis, dissimilatory nitrate reduction, histidine degradation, fumarate reductase (prokaryotes), sulfate-sulfur assimilation, hydroxyproline degradation, phylloquinone biosynthesis, C1-unit interconversion (prokaryotes), cytochrome-c oxidase (cbb3-type), phosphatidylethanolamine biosynthesis, D-galacturonate degradation (bacteria), cytochrome-c oxidase (prokaryotes), and cytochrome bc1 complex respiratory subunit (Fig. 6b). Acetogenesis, the process of producing acetate via the reduction of CO_2_ anaerobically via the Wood–Ljungdahl pathway, is not a known functional pathway of bacteria identified in this study. Upon further inspection, two key enzymes are present: acetate kinase (AckA) and phosphotransacetylase (Pta). These two proteins are the main enzymes of the PTA-ACK pathway, which mediates the conversion of acetyl-CoA to acetate by using acetyl phosphate; acetate is either consumed or excreted depending on cellular demand [110]. Cobalamin (vitamin B12) biosynthesis was another module that decreased post-induction; however, only the cobyrinate a,c-diamide => cobalamin conversion was significant. *Celeribacter halophilus* (CT-PreInd-bin.7) was the only bMAG that encoded genes for additional cobalamin biosynthesis steps (uroporphyrinogen III => sirohydrochlorin => cobyrinate a,c-diamide), and its relative abundance was not impacted by induction.

**Figure 6:**
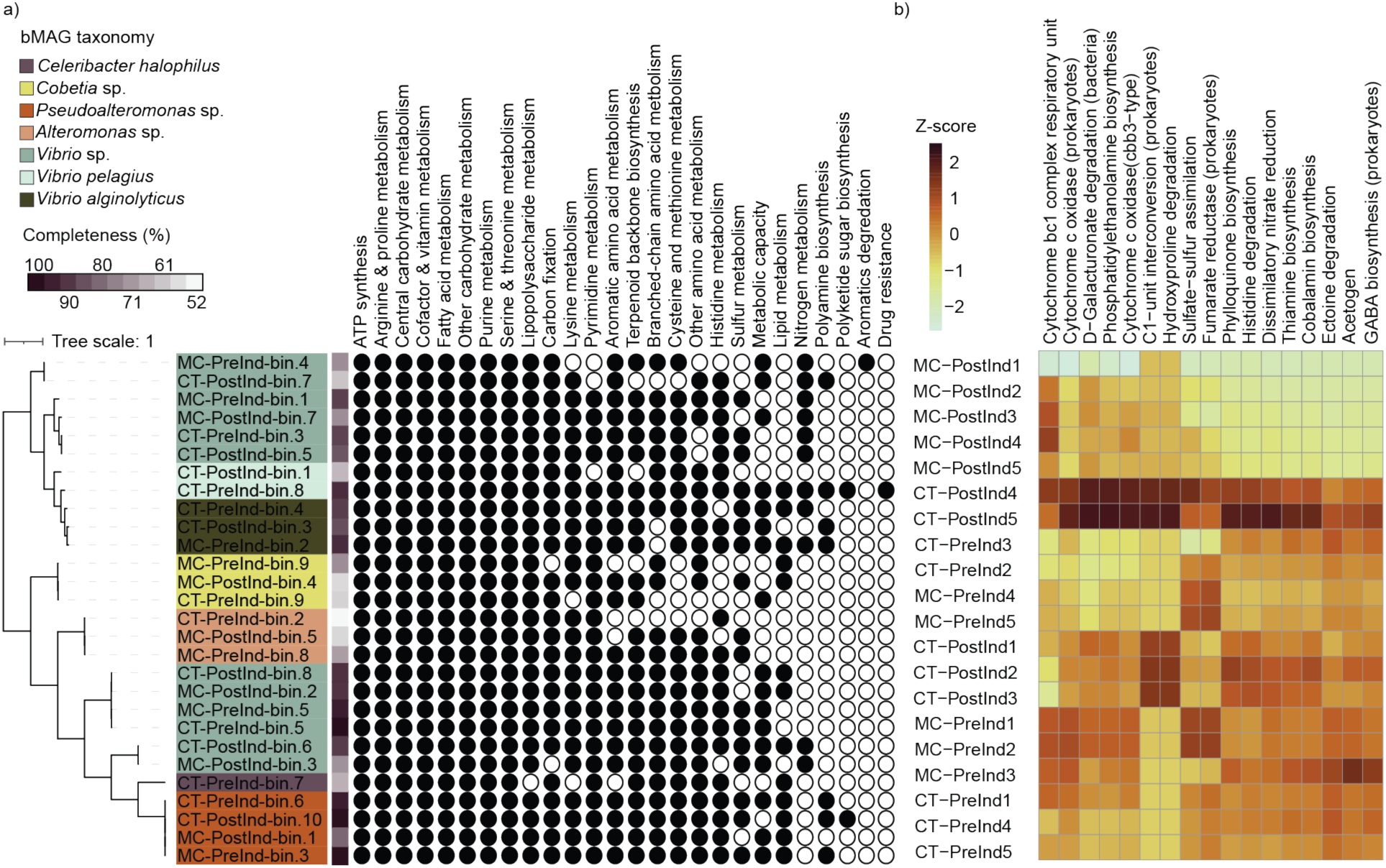
Bacterial metabolism profile is altered post-induction. (a) Average Nucleotide Identity (ANI) tree for all 28 identified bMAGs. bMAG completeness is represented by the dark purple to white color gradient, and taxonomy corresponds to the color of the tree node. Black circles indicate KEGG metabolism submodules that have at least one module with a pathwise completion greater than 70%. (b) KEGG metabolism modules that SIMPER (*p* < 0.05) identified as driving differences between the MC and control post-induction samples. The relative abundance of each bMAG that encoded the module was summed within a replicate and was normalized by module with a Z-score.

Although their pathwise completion was less than 70%, genes for six different pathogenicity modules with pathwise completion rates of 4-33% were present. ‘*Vibrio cholerae* pathogenicity signature, cholera toxins’ genes were identified in seven *Vibrio* bMAGs and ranged from 16-33% pathwise completion. Two *Vibrio* sp. and one *Vibrio pelagius* bMAGs encoded zonula occludens toxin (Zot), which is a prophage-encoded enterotoxin that increases the virulence of *Vibrio* species [111], and one *Vibrio pelagius* and two *Vibrio alginolyticus* bMAGs encoded thermolabile hemolysin (*tlh*) (Data S10). One *Vibrio alginolyticus* bMAG encoded both *zot* and *tlh*.

## Discussion

Biofilm formation is a common capacity of eukaryotic microbiomes, and the ability to grow as biofilms often regulates community composition [112–114]. Over 60% of the bacterial and viral community members in the *in vitro* multispecies biofilms described here were also present and abundant in *in situ* samples of *S. natans* from South Florida (Fig. 1d and 1e), including several *Vibrio* spp. and *Vibrio* prophages. These results demonstrate that our experiments involve ecologically relevant taxa. Several mechanisms regulate biofilm growth, and prophage induction can have significant impacts on biofilm development and the functioning of the eukaryotic host [115]. For example, a filamentous prophage in *Pseudomonas aeruginosa* strain PAO1 contributes to biofilm development through host autolysis, creating hollow centers within the biofilm that facilitate cell dispersal and ultimately promote biofilm growth [116, 117]. In marine biofilms, the impacts of prophages remain mostly unknown, despite their widespread presence and abundance [118–120]. Some of the known examples include the prophage-influenced niche switch from planktonic to biofilm growth in *Sulfitobacter* sp. strain CB2047 [49], isolated from microalgae. Prophage induction has also been shown to cause decreased cell growth, increased copies of phage tail genes, and electroactive biofilm decay in *Geobacter sulfurreducens* biofilms [121]. Conversely, several studies in non-marine systems have demonstrated that cell lysis via spontaneous prophage induction enhances biofilm formation of single species biofilms by releasing extracellular DNA and other components necessary for biofilm production [122–125]. The data described here shows that the prophage induction of a *Sargassum*-derived multispecies biofilm by MC resulted in a decrease in biofilm formation 12 hours after the induction (Fig. 1c) [35], which may involve cell toxicity by the inducting agent in addition to viral lysis. Therefore, understanding the impact of prophage induction, spontaneous or via an exogenous agent, is crucial for understanding biofilm dynamics in the marine environment.

### Prophage induction modulates bacterial community structure

Among the 11 *Sargassum*-derived prophages identified in this dataset, seven were predicted to infect bacteria in the genus *Vibrio*, including four prophages that drove most of the differences in viral community composition between the control and MC post-induction groups (Fig. 2b). These results suggest that in *Vibrio*-dominated *Sargassum* microbiomes, as is the case for most Caribbean and Florida samples [35, 36, 38], phage integration and induction may have a significant impact on community composition and dynamics. Prophage induction with MC has been shown to introduce microbiome shifts in other systems, such as soil, resulting in an increase in bacterial diversity within 24 hours [126], which could also be the case in this system. *Vibrio*-infecting prophages identified here were classified as *Corticoviridae,* a group of tailless double-stranded DNA phages present in the majority of marine *Vibrio* genomes [127], and tailed double-stranded DNA phages in the *Caudoviricetes* group. One *Inoviridae* phage, a filamentous virus also common in *Vibrio*s [127], displayed an increase in abundance in the MC-treated group (Fig. 3c), but could not be confirmed as a prophage due to the lack of bacterial flanking regions or an episomal replication style [128]. Combined with the negative change in relative abundance of *Vibrio* host bMAGs in the MC post-induction group (Fig. 4b), the presence of these broadly distributed prophage groups in our inducible dataset from *Sargassum* further supports the idea of phage induction of the genus *Vibrio* being a potential mechanism controlling *Vibrio* abundances in *Sargassum*. A previous induction study showed that *Sargassum* prophages can be induced not only with MC, but also by ultraviolet light [35], albeit to a lesser extent. Therefore, prophage induction may be widespread in naturally occurring *Sargassum* with pronounced effects when *Vibrio* is the dominant group.

The decrease in *Vibrio* bMAG abundance due to induction was accompanied by an increase in the relative abundance of two *Pseudoalteromonas* sp. and one *Cobetia* sp. (and one *Vibrio* sp. to a lesser extent; Fig. 4b) and the persistence of one *Alteromonas* sp. that displayed a decrease in relative abundance in the control. This suggests that in the short term, prophage induction decreases *Vibrio* spp. abundance while allowing other bacteria to persist or start growing. Metataxonomic studies of *Sargassum* microbiome composition across its range have reported that when *Gammaproteobacteria* comprise the majority of the *Sargassum* microbiome, there are two distinct profile compositions: one with a greater proportion of *Vibrio* and another with a greater proportion of *Alteromonas* [37]. Our induction data is consistent with and may help explain these observations. Further, prophage induction has been documented as a mechanism for killing bacterial competitors [129, 130]. The coral pathogen *Vibrio coralliilyticus* induces a prophage in a nonpathogenic *Vibrio* sp. through hydrogen peroxide production, killing the commensal *Vibrio* sp. and allowing *Vibrio coralliilyticus* to colonize coral tissue [130].

Therefore, mass prophage induction may be a biotic factor impacting the alternating taxonomic dominance in *Sargassum* microbiomes across its geographical range. Another factor contributing to these community composition shifts is the senescence of the algae, as evidenced by stranding simulations with *Sargassum,* which demonstrate that algae decay favors *Vibrio* [131]. Future studies analyzing prophage induction during senescence will be able to test whether prophage induction is reduced during algae decay.

### Prophages impact on quorum sensing, virulence, and biofilm formation

Quorum sensing (QS), bacterial cell-to-cell communication, mediates individual and population behaviors and is controlled by extracellular signaling molecules known as autoinducers [132]. In *Vibrio*, two master regulators control quorum sensing: AphA, a part of the PadR protein family, which operates at low cell densities, and the LuxR protein family, which operates at high cell densities [87]. These two transcriptional regulators control the expression of one another and a vast array of other genes [133, 134]. Here, we identified a *Vibrio* prophage-encoded version of AphA, the low cell density master regulator of *Vibrio* QS. At low cell densities, bacterial AphA is upregulated, leading to the activation of biofilm formation and virulence genes, thereby enhancing colonization [87, 106, 107, 133–135]. The prophage-encoded *aphA* gene was flanked by viral genes and shared only 34% amino acid similarity with the host homolog, indicating it was unlikely to be a result of host contamination. Despite this sequence divergence, the two proteins are structurally similar (Fig. 5b). The prophage-encoded AphA is most closely related to another viral-encoded protein from a phage identified in *S. natans* metagenomes. Both viral proteins were closely related to a non-redundant version of AphA found in multiple *Vibrio* species (Fig. 5a), suggesting that the prophage acquired it from a previous *Vibrio* host. QS state of *Vibrio anguillarum* has been demonstrated to mediate its H20-like prophage’s lysogeny decisions, and this prophage stimulates biofilm growth at low cell densities [136]. Analogous to phage-encoded LuxR-type transcription regulators, which allow phages to eavesdrop on their bacterial host’s quorum-sensing network to inform their lysis-lysogeny decisions, including hosts in *Vibrio* and *Aeromonas* [137, 138], prophage-encoded AphA may similarly influence the lysogenic switch or bacterial gene regulation. Conversely, the induction of this prophage may decrease biofilm formation capacity in the *Vibrio* host.

*Vibrio* temperate phages give fitness advantages and disadvantages to their host. One benefit is that prophage-encoded toxins enhance host virulence. A classic example is the cholera toxin-encoding filamentous temperate phage CTXΦ of *Vibrio cholera* [139]. Other examples of *Vibrio* phage-encoded toxins are RTX and Zot [139, 140]. The C-terminal end of the Zot protein is involved in phage assembly, but the N-terminus is cleaved and secreted, where it breaks down tight junctions [141]. Here, we identified one *Inoviridae* phage, encoding Zot, whose increase in fractional abundance in the MC-treated group was partially responsible for the differences between treatments. Macroalgal cells do not form tight junctions; instead, they form plasmodesmata for intercellular communication and adhesion [142]. Therefore, Zot may be used in the infection of animals associated with *Sargassum* or may be involved in other processes mediated by actin filaments in algal cells.

The temperate phage that showed the second largest increase in abundance in the MC-treated group encoded a putative chitinase gene (Fig. 3c and 3d)*. Vibrio* often associate with organisms that produce chitin, such as crustaceans, zooplankton, bivalves, and even some bryozoans [143, 144], and possess a conserved set of genes for chitin degradation, facilitating its utilization as a carbon and nitrogen source [145]. Although *Sargassum* itself does not produce chitin, holobiont members associated with *Sargassum*, like bryozoans, incorporate chitin and calcium carbonate into their exoskeletons [146, 147]. *Vibrio* spp. chitinases have been identified in several *Vibrio* infecting prophages [148], and a *Pseudomonas* sp. prophage encoding a chitinase enhanced the ability of its host, isolated from deep-sea sediments, to degrade chitin and form biofilms [48]. The identification of prophage chitinases in many *Vibrio* bacteria suggests lysogenic conversion of chitinases could be common in other marine environments.

### Prophage induction alters the metabolic profiles of Sargassum biofilms

Seventeen metabolic modules encoded by the bacterial community differed between post-induction groups, all of them decreasing in the MC-treated group (Fig. 6b). Among the metabolisms depleted by induction was dissimilatory nitrate reduction (Fig. 6b). Nitrogen metabolism is of interest concerning *Sargassum* because the algae’s tissue % N has increased by 35% since the 1980s, demonstrating that increased N availability is a factor contributing to *Sargassum* blooms [149]. *Sargassum* δ^15^N, an indicator of N source, varies across its range, indicating that the algae utilize multiple N pools [149, 150]. In certain regions, low δ^15^N and nitrogen fixation rates of *Sargassum* mats signal that N fixation by bacterial epiphytes on *Sargassum* is contributing to the N sources of these blooms [32, 33, 149, 150]. A study examining the relative abundance of genes involved in nitrogen cycling, using *nifH* as an indicator of nitrogen fixation, revealed that the majority of nitrogen-fixing bacteria on *Sargassum* were heterotrophic bacteria in the genera *Vibrio* and *Filomicrobium* [32]. In our study, we identified one *Vibrio* sp. bMAG that encoded the *nifH* enzyme; this bacterium also had genes for dissimilatory nitrate reduction and nitrate assimilation (Data S10). Several *Vibrio* spp. are documented as nitrogen fixers [151, 152]. ^15^N isotope tracing studies with sea urchins have demonstrated that microbially fixed N, via a *Vibrio* bacterium, can be incorporated into sea urchin tissues [153], suggesting that *Vibrio* bacteria could supply *Sargassum* with fixed N. Dissimilatory nitrate-reducing bacteria decreased upon prophage induction, whereas nitrate assimilation was not impacted. Overall, these findings suggest that prophage induction has the capacity to shape key N recycling microbial abundances in the *Sargassum* microbiome.

## Conclusion

Our study shows that prophage induction can significantly alter the community composition and metabolic capacity of multispecies biofilms representative of heterotrophic bacteria associated with pelagic *Sargassum*. Prophage induction reduced *Vibrio* abundances and opened niche space for less abundant *Gammaproteobacteria* in the genera *Alteromonas*, *Pseudoalteromonas*, and *Cobetia*. This reduction in *Vibrio* bacteria has the capacity to alter sulfur, nitrogen, and carbon metabolisms within the biofilm community. Induced viruses encoded genes involved in biofilm formation, virulence, and host metabolism. Overall, the data from this study suggest that viral infection strategy may be a biotic factor influencing *Vibrio* abundances associated with *Sargassum*, providing a potential mechanism that contributes to changes in community composition across *Sargassum*’s range.

## Author contributions

CBS and AKS designed the study. AKS, NSV, and BAW analyzed the genomic data. AKS analyzed the results and wrote the manuscript. All authors contributed to the manuscript review process.

## Acknowledgments

We thank the Frost Institute for Data Science and Computing (IDSC) for providing access to the University of Miami’s high-performance computing system.

## Data availability

Raw reads are available in the NCBI Sequence Read Archive (PRJNA1111456 for metagenomes from biofilm induction assay [Figure 2-6] and PRJNA1063307 for metagenomes of naturally occurring *S. natans* VIII [Figure 1d & 1e]). Co-assembled contigs [154], bMAGs and the Newick file for the ANI tree [155], prophages with and without flanking regions [156], biofilm assay viral sequences [157], and AphA protein sequences and the Newick file for their protein phylogeny tree [158], are available on Figshare under the [Sargassum-derived biofilm induction project] project, [https://figshare.com/projects/Sargassum-derived_biofilm_induction_project/260945]. The code used for the bioinformatic pipeline and the R script for data analysis are available on the Silveira-Lab GitHub under the [Sargassum-derived-Biofilm-Induction-Bioinformatics-Pipeline] repository, [https://github.com/Silveira-Lab/Sargassum-derived-Biofilm-Induction-Bioinformatics-Pipeline].

## Supplementary information

Supplementary figures and tables: Fig. S1-5 and Table S1-5. Figure and table legends are included in the attached document.

Data S1: Biofilm induction experiment bMAG GTDB-tk taxonomy, completeness, contamination, completeness model, genome size, total contigs, GC content, and relative abundance in genomes per million reads per replicate (columns I-AB).

Data S2: Biofilm induction experiment non-dereplicated viral sequences (*n* = 1,304) GC content, length, CDS count, viral population number, and whether that viral sequence is the population representative.

Data S3: Biofilm induction experiment dereplicated, quality-filtered viral fractional abundances in the *in situ S. natans* metagenome sampled and collected off the coast in South Florida. The datasheet contains the viral genome ID, *S. natans* sample ID, number of reads mapped to viral genome, coverage percent, the number of all biofilm induction virus mapped reads, fractional abundance, infection strategy prediction and probability, if the viral genome is a prophage, viral realm, phylum, class, order, and family.

Data S4: Biofilm induction experiment bMAG relative abundance, in genomes per million reads, in in situ *S. natans* and seawater samples collected off the coast of South Florida. The datasheet contains the genome bin ID, GTDB-tk taxonomy, and the relative abundance of each genomic bin in *S. natans* and seawater replicates.

Data S5: RefSeq GCF genome BLASTn to bacterial flanking regions associated with prophages. This datasheet contains the bacterial flank IDs, RefSeq GCF genome accession numbers, the genus of the RefSeq GCF genome, percent identity, length of the bacterial flank, length of the RefSeq GCF genome, match length, number of mismatches, the start and end positions of the match for both the bacterial flank and the RefSeq GCF genome, and the e-value.

Data S6: Virus-to-host ratios in biofilm induction. This datasheet contains the prophage viral genome ID, shortened prophage ID, replicate, sample type, prophage-to-host ratio (VHR), prophage mean and median coverage, prophage coverage standard deviation, prophage coverage breadth, host mean and median coverage, and host coverage standard deviation.

Data S7: Gene annotations for viruses in Fig. 2c and Fig. 3d. The datasheet contains the viral gene ID, the viral ID, gene start, gene end, strand, gene functional category, MetaCerberus annotation, the database match, and the e-value.

Data S8: Biofilm induction experiment virus categorical and fractional abundance data. The datasheet contains contig ID, viral ID, virus length, number of reads mapped to the virus, the number of reads mapped to all biofilm induction viruses, the average contig length, fractional abundance, infection strategy, infection strategy probability, if the virus is a prophage, viral realm, phylum, class, order, and family.

Data S9: AphA BLASTp matches against the non-redundant RefSeq protein database. The datasheet contains the virus or host ID, the AphA protein ID from this study, the accession number of RefSeq protein matches, percent identity, coverage of the AphA proteins from this study, protein lengths, match length, number of mismatches, the start and end position of the matches, and the e-value.

Data S10: Biofilm induction experiment bMAG metabolism data. This datasheet contains the KEGG module ID, bMAG ID, module name, module class, module category, module subcategory, module definition, stepwise module completeness, pathwise module completeness, proportion of unique enzymes present, KEGG IDs of enzymes unique to the module, number of unique enzymes found, enzyme hits in the module, and warnings for interpretation.

