## Supplementary figures and tables for "Prophage induction shifts community composition and functional capacity in a *Sargassum*-derived multispecies biofilm"

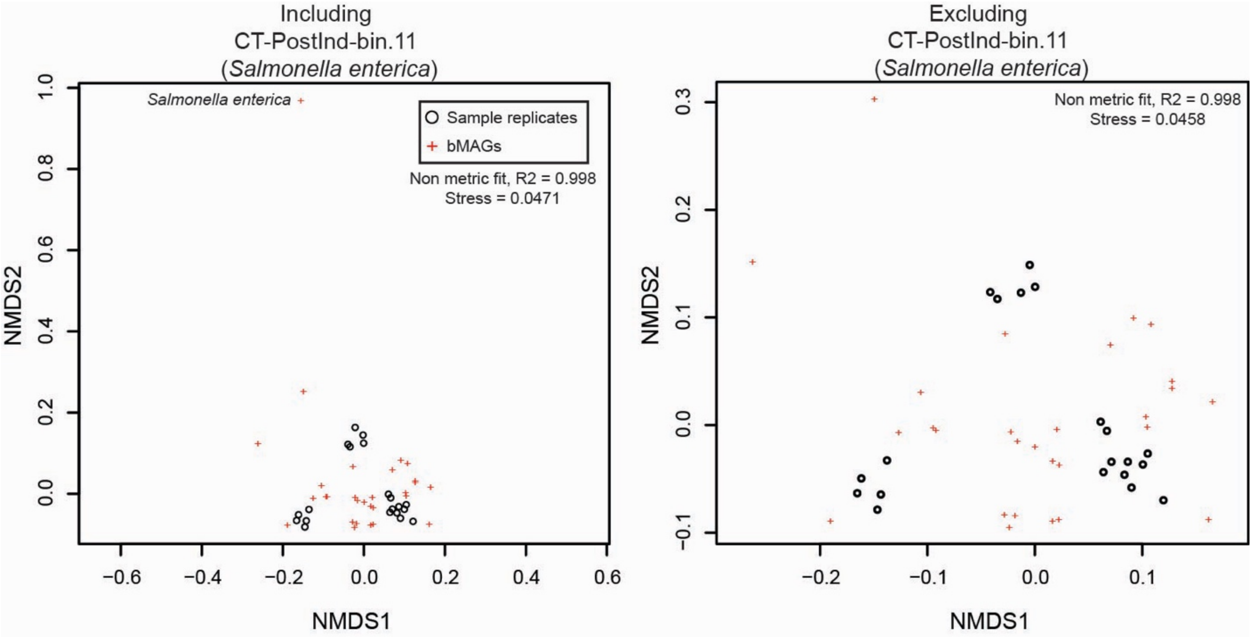


**Fig S1: *Salmonella enterica* bMAG contaminant is an outlier** | a) NMDS plot comparing the bMAG relative abundance in genomes per million reads per replicate with *Salmonella enterica* contaminant included. Black circles and red crosses represent sample replicates and bMAGs, respectively. b) NMDS plot comparing the bMAG relative abundance in genomes per million reads per replicate with *Salmonella enterica* contaminant removed.

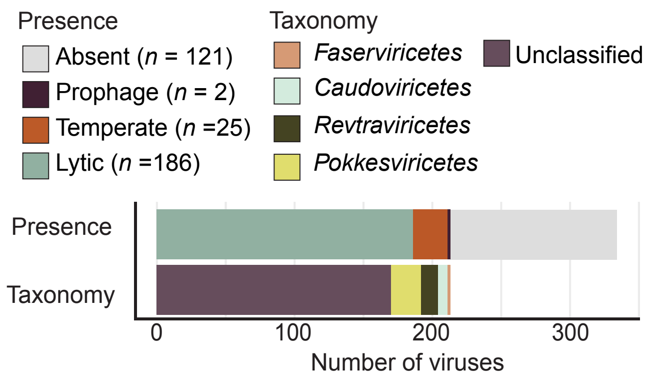


**Fig. S2: Presence of biofilm assay viruses in environmental *S. natans* metagenomes |** Presence of the 334 viruses identified in the biofilm induction experiment in *in situ S. natans* samples, with colors representing absent viruses (gray), present prophages (dark purple), temperate (orange), and lytic viruses (blue). Taxonomy of the 213 viruses present in *S. natans* samples: *Faserviricetes* (light orange), *Caudoviricetes* (light blue), *Revtraviricetes* (dark green), and *Pokkesviricetes* (light green).

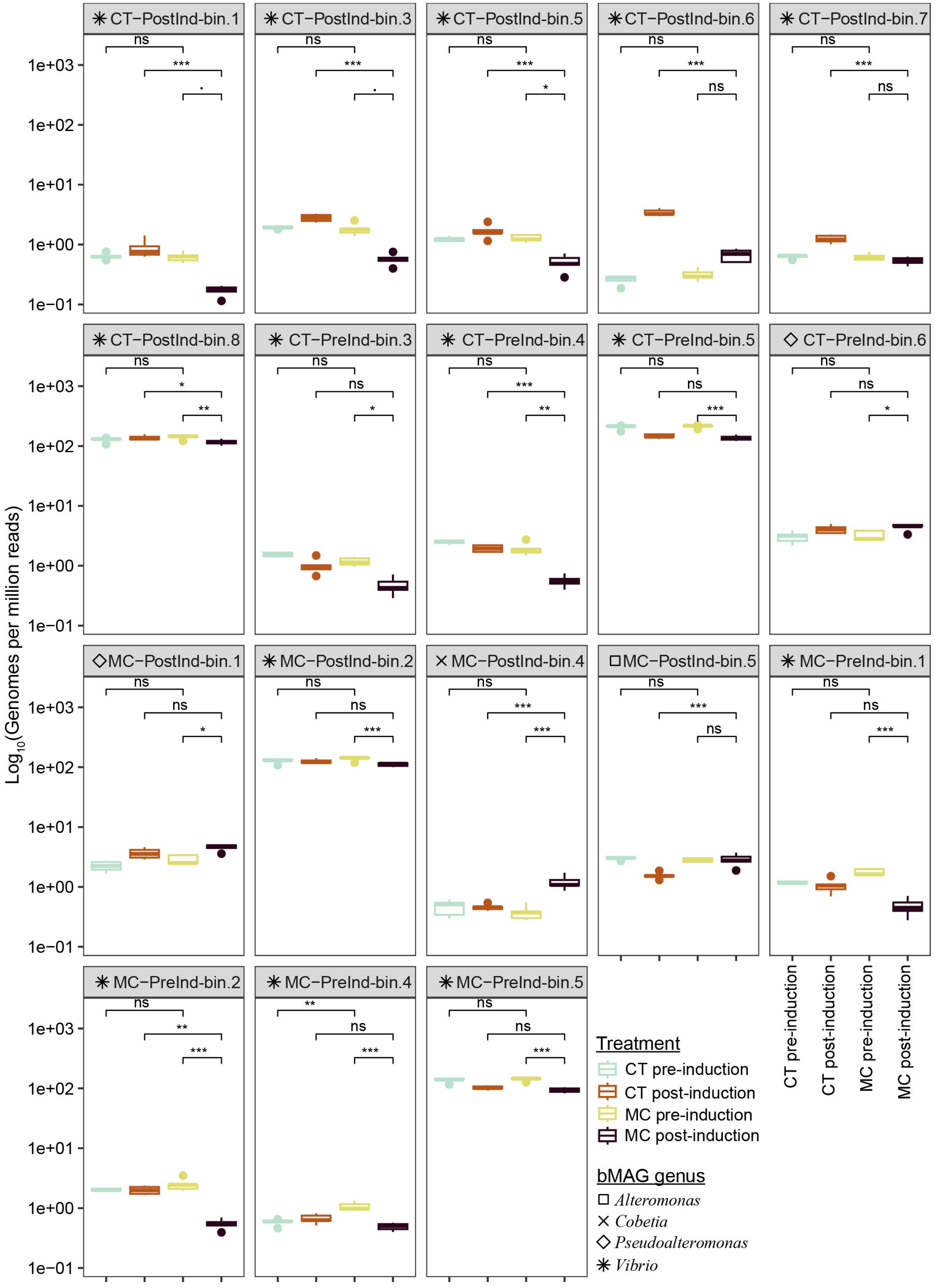

CT−PostInd-bin.1
CT−PostInd-bin.3
CT−PostInd-bin.5
CT−PostInd-bin.6
CT−PostInd-bin.7
CT−PostInd-bin.8
CT−PreInd-bin.3
CT−PreInd-bin.4
CT−PreInd-bin.5
CT−PreInd-bin.6
MC−PostInd-bin.1
MC−PostInd-bin.2
MC−PostInd-bin.4
MC−PostInd-bin.5
MC−PreInd-bin.1
MC−PreInd-bin.2
MC−PreInd-bin.4
MC−PreInd-bin.5
Log10(Genomes per million reads)
1e+03
1e+02
1e+01
1e+00
1e−01
CT pre-induction
CT post-induction
MC pre-induction
MC post-induction
Treatment
bMAG genus
Alteromonas
Cobetia
Pseudoalteromonas
Vibrio

**Fig. S3: 18 bMAGs are differentially abundant when comparing control and Mitomycin C post-induction groups and Mitomycin C pre- and post-induction groups** | Boxplots for the 18 bMAGs identified as driving differences between control and MC post-induction groups and MC pre- and post-induction groups. Each group of boxplots represents an individual bMAG, labeled in the gray bar above the plot. The genus of each bMAG is depicted with shapes: *Alteromonas* (square), *Cobetia* (x), *Pseudoalteromonas* (diamond), and *Vibrio* (star). The x-axis represents sample types, which are also color-coded with CT pre-induction, CT post-induction, MC pre-induction, and MC post-induction being light blue, orange, light green, and dark purple, respectively. SIMPER *p*-values are shown with asterisks (*** $p$-value < 0.001, ** $p$-value < 0.01, * $p$-value < 0.05, ns $p$-value > 0.05).

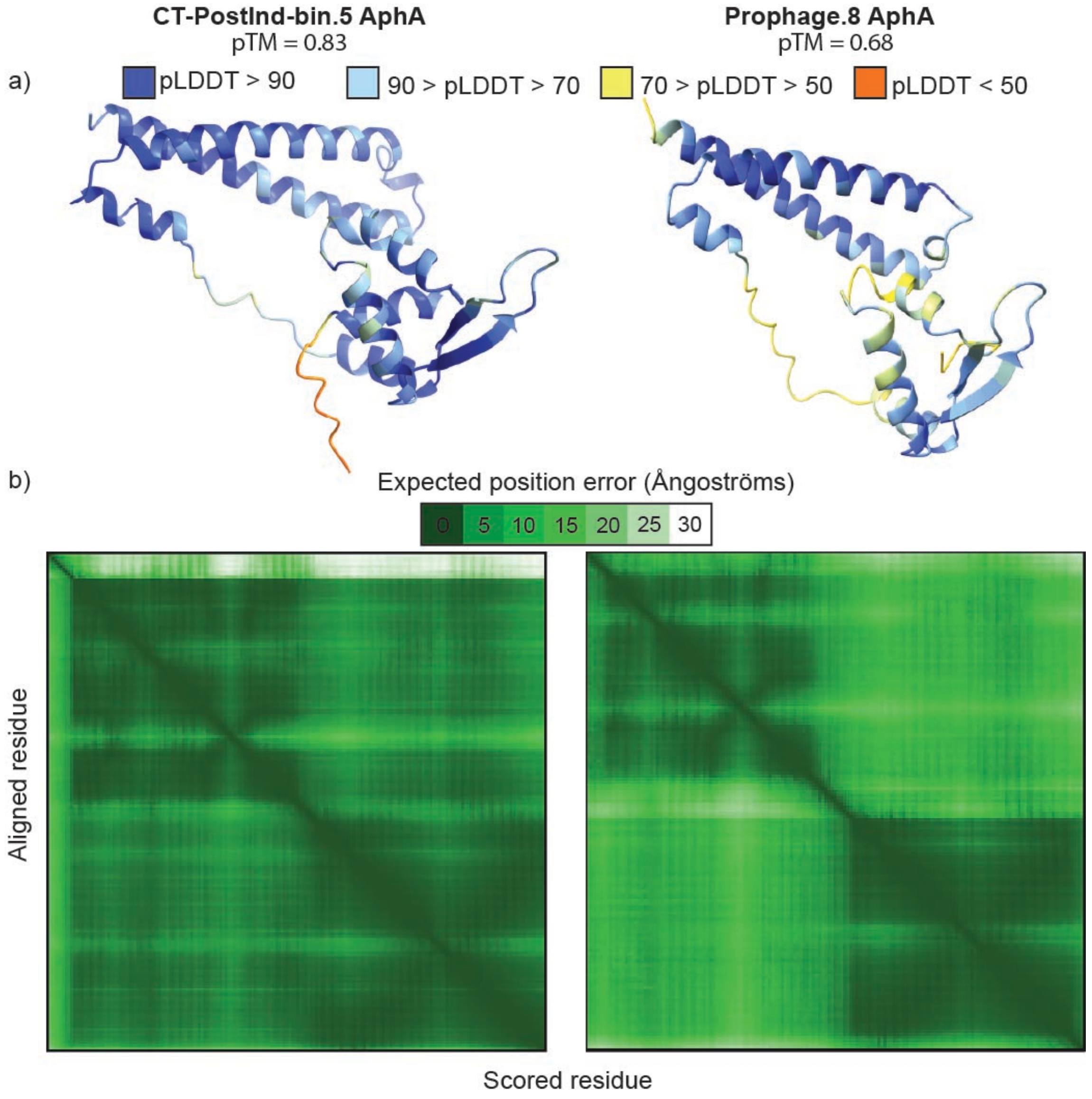


**Fig. S4: Prophage.8 and host-encoded AphA protein structures** | a) Alphafold3 predicted structures for prophage.8 (right) and its *Vibrio* sp. host (left). Colors represent a per-atom confidence estimate (pLDDT) on a 0-100 scale; higher values (dark and light blue) indicate a higher confidence. Predicted template modeling (pTM) scores are listed below the protein's name. b) Predicted aligned error (PAE) plots for each protein, which estimate the error in relative position between two tokens in the predicted structure. Lower PAE values (darker green) indicate a higher confidence.

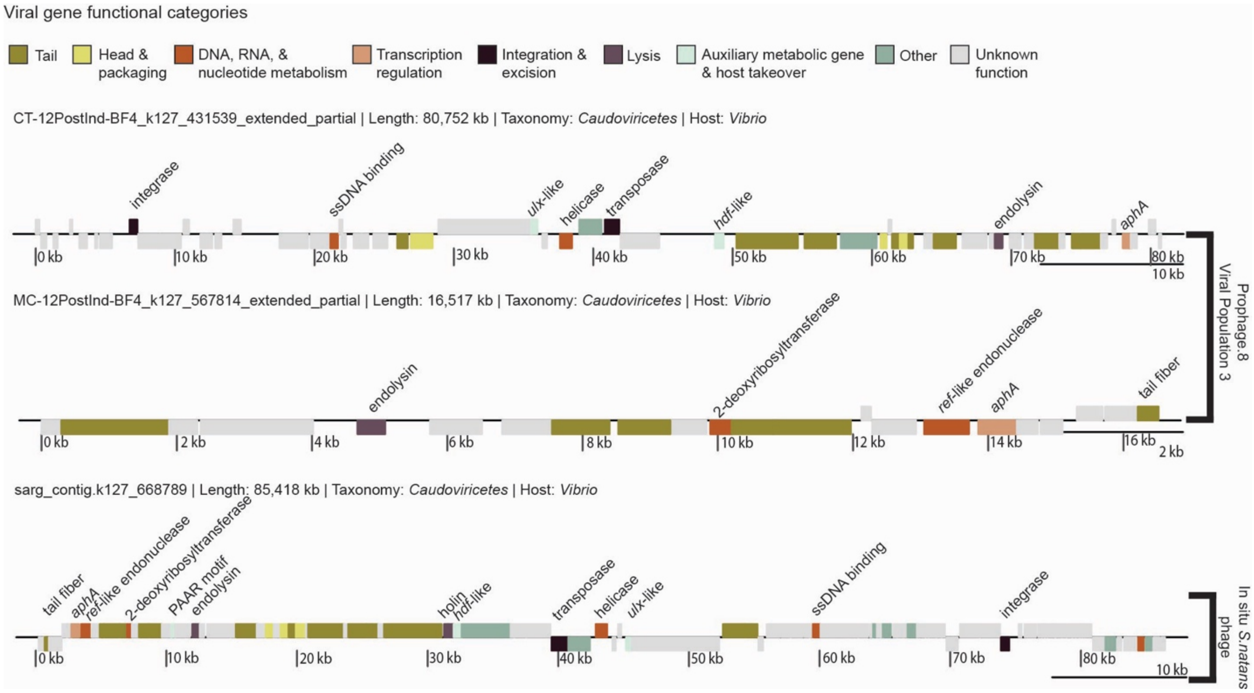


**Fig. S5: The gene encoding the AphA protein is flanked by viral genes** | Genome plots of three phages that encode the *aphA* gene. The first two phages are the population representative (k127_431539) for prophage.8 and the viral sequence that was classified as a prophage with the bacterial flanking regions removed (k127_567814). The third plot is a phage (k127_668789) from in situ *S. natans* metagenome samples collected in South Florida in March of 2021.

### Supplementary Tables

**Table S1: Raw read file information** | Read file names, barcode sequences, number of raw reads, yield in Megabases, mean quality score, percent of bases greater than or equal to 30, and the number of reads after quality control.

| Sample ID | Barcode Sequence | # Raw Reads | Yield (Mbases) | Mean Quality Score | % Bases >= 30 | # QC Reads |
|---|---|---|---|---|---|---|
| CT-12PostInd-BF1 | CGCTCATT+GTCAGTAC | 31365608 | 9410 | 39.14 | 95.72 | 31018570 |
| CT-12PostInd-BF2 | GAGATTCC+AGGCTATA | 28605902 | 8581 | 39.13 | 95.68 | 28245572 |
| CT-12PostInd-BF3 | GAGATTCC+GCCTCTAT | 28332429 | 8499 | 39.16 | 95.76 | 27992969 |
| CT-12PostInd-BF4 | GAGATTCC+AGGATAGG | 31366036 | 9410 | 39.08 | 95.42 | 30940900 |
| CT-12PostInd-BF5 | GAGATTCC+TCAGAGCC | 33030845 | 9910 | 39.05 | 95.25 | 32564589 |
| CT-PreInd-BF1 | TCCGGAGA+TAAGATTA | 34365729 | 10310 | 38.92 | 94.63 | 33825707 |
| CT-PreInd-BF2 | TCCGGAGA+ACGTCCTG | 32954812 | 9886 | 39.17 | 95.85 | 32584548 |
| CT-PreInd-BF3 | TCCGGAGA+GTCAGTAC | 32796949 | 9839 | 38.98 | 94.92 | 32336597 |
| CT-PreInd-BF4 | CGCTCATT+AGGCTATA | 32241223 | 9672 | 39.09 | 95.46 | 31846724 |
| CT-PreInd-BF5 | CGCTCATT+GCCTCTAT | 35288042 | 10586 | 39.1 | 95.49 | 34854390 |
| MC-12PostInd-BF1 | GAGATTCC+CTTCGCCT | 33697573 | 10109 | 39.12 | 95.59 | 33291234 |
| MC-12PostInd-BF2 | GAGATTCC+TAAGATTA | 32673448 | 9802 | 39.1 | 95.5 | 32240691 |
| MC-12PostInd-BF3 | GAGATTCC+ACGTCCTG | 30165351 | 9050 | 39.15 | 95.75 | 29801776 |
| MC-12PostInd-BF4 | GAGATTCC+GTCAGTAC | 30584124 | 9175 | 38.81 | 94.41 | 30224016 |
| MC-12PostInd-BF5 | ATTCAGAA+AGGCTATA | 30480550 | 9145 | 39.08 | 95.41 | 30085614 |
| MC-PreInd-BF1 | CGCTCATT+AGGATAGG | 31104206 | 9331 | 39 | 95.02 | 30658324 |
| MC-PreInd-BF2 | CGCTCATT+TCAGAGCC | 31042080 | 9313 | 38.94 | 94.72 | 30560354 |
| MC-PreInd-BF3 | CGCTCATT+CTTCGCCT | 36344338 | 10903 | 39.12 | 95.59 | 35888184 |
| MC-PreInd-BF4 | CGCTCATT+TAAGATTA | 30769127 | 9230 | 39.04 | 95.2 | 30350760 |
| MC-PreInd-BF5 | CGCTCATT+ACGTCCTG | 32667869 | 9800 | 39.01 | 95.05 | 32212742 |

**Table S2: BLASTn results of Prophage.8 and its bacterial flanking region against a virus binned to a *Vibrio* bMAG without identified flanking regions** | Sequence ID, query ID, percent identity, sequence length, query length, match length, and sequence and query start and end positions for the BLASTn comparison of prophage.8 and its bacterial flanking region to viral sequence (CT-PostInd-k127_431539) binned to a *Vibrio* bMAG (CT-PostInd-bin.5).

| Sequence ID | Query ID | % ID | Sequence length | Query length | Match length | Sequence start | Sequence end | Query start | Query end | e-value |
|---|---|---|---|---|---|---|---|---|---|---|
| **Prophage.8** | CT-PostInd-k127_431539 | 99.927 | 16517 | 81307 | 16517 | 1 | 16517 | 64143 | 80659 | 0.0 |
| **Prophage.8 flank** | CT-PostInd-k127_431539 | 99.383 | 3166 | 81307 | 648 | 1 | 648 | 80660 | 81307 | 0.0 |

**Tables S3: Prophage identification, taxonomy, host, bacterial flank, and coordinates information** | General information for all 11 identified prophages, including their shortened prophage ID, identification method, taxonomy, iPHoP predicted host (more info in Table S4) , if the prophage had a bMAG link, bacterial flank BLASTn taxonomy (BLASTn results in Data S5), length, prophage start residue in original contig, prophage end residue in original contig, length of bacterial flanks, viral genome ID, and the viral population that prophage was assigned to.

| Prophage ID | ID method | Taxonomy | IPHoP host | bMAG match | GTDB BLASTn | Prophage length | start | end | Flank 1 length | Flank 2 length |
|---|---|---|---|---|---|---|---|---|---|---|
| prophage.1 | Genomad | Caudovircetes | NA | NA | NA | 29570 | 16859 | 46428 | 16858 | 379 |
| prophage.2 | CheckV | Caudoviricetes | Rickettsia | NA | Streptomyces | 7111 | 11881 | 18991 | 11880 | 0 |
| prophage.3 | Genomad | Corticoviridae | NA | CT-PostInd-bin.8 | Vibrio | 7548 | 3704 | 11252 | 3703 | 2 |
| prophage.4 | CheckV | Caudoviricetes | NA | NA | NA | 1375 | 5775 | 7149 | 5774 | 0 |
| prophage.5 | CheckV, Genomad | Caudoviricetes | Vibrio | NA | NA | 29343 | 2 | 29344 | 1 | 33273 |
| prophage.6 | CheckV | Kyanoviridae | NA | NA | Vibrio | 1321 | 2601 | 3921 | 2600 | 0 |
| prophage.7 | CheckV, Genomad | Caudoviricetes | NA | NA | Vibrio | 119072 | 16409 | 135480 | 16408 | 40224 |
| prophage.8 | Genomad | Caudoviricetes | Vibrio | CT-PostInd-bin.5 | Vibrio | 16517 | 3 | 16519 | 2 | 3166 |
| prophage.9 | CheckV | Caudoviricetes | Vibrio | MC-PreInd-bin.2 | Vibrio | 32400 | 7906 | 40305 | 7905 | 0 |
| prophage.10 | Genomad | Autographiviridae | Vibrio | NA | NA | 9075 | 2183 | 11257 | 2182 | 0 |
| prophage.11 | Genomad | Caudoviricetes | NA | NA | NA | 40302 | 2 | 40303 | 1 | 6632 |

**Table S3 (continued).**

<table>
<tr><th>Prophage ID</th><th>Viral genome ID</th><th>Viral population rep ID</th></tr>
<tr><td>prophage.1</td><td>CT-12PostInd-BF1_megahit_k127_173951_flag_0_multi_10693.0828_len_47076|provirus_16859_46428</td><td>CT-12PostInd-BF1_megahit_k127_173951_flag_0_multi_10693.0828_len_47076|provirus_16859_46428</td></tr>
<tr><td>prophage.2</td><td>CT-12PostInd-BF1_megahit_k127_412173_flag_1_multi_6.0000_len_18991</td><td>CT-12PostInd-BF1_megahit_k127_412173_flag_1_multi_6.0000_len_18991_1</td></tr>
<tr><td>prophage.3</td><td>CT-12PostInd-BF2_megahit_k127_162855_flag_0_multi_236.2276_len_8955_extended_partial|provirus_3704_11251</td><td>MC-12PostInd-BF1_megahit_k127_555745_flag_0_multi_68.4010_len_7549_extended_partial</td></tr>
<tr><td>prophage.4</td><td>CT-PreInd-BF1_megahit_k127_212264_flag_0_multi_12.2817_len_7149</td><td>CT-PreInd-BF1_megahit_k127_212264_flag_0_multi_12.2817_len_7149_1</td></tr>
<tr><td>prophage.5</td><td>CT-PreInd-BF1_megahit_k127_349326_flag_1_multi_11.9815_len_62618|provirus_2_29344</td><td>CT-PreInd-BF1_megahit_k127_349326_flag_1_multi_11.9815_len_62618|provirus_2_29344</td></tr>
<tr><td>prophage.6</td><td>CT-PreInd-BF1_megahit_k127_714724_flag_1_multi_5.9987_len_3921</td><td>CT-PreInd-BF1_megahit_k127_714724_flag_1_multi_5.9987_len_3921_1</td></tr>
<tr><td>prophage.7</td><td>MC-12PostInd-BF1_megahit_k127_435789_flag_0_multi_7452.9043_len_35695_extended_partial|provirus_16409_135480</td><td>MC-12PostInd-BF1_megahit_k127_435789_flag_0_multi_7452.9043_len_35695_extended_partial</td></tr>
<tr><td>prophage.8</td><td>MC-12PostInd-BF1_megahit_k127_567814_flag_0_multi_29.0000_len_13848_extended_partial|provirus_3_16519</td><td>CT-12PostInd-BF4_megahit_k127_431539_flag_0_multi_14.0000_len_69873_extended_partial</td></tr>
<tr><td>prophage.9</td><td>MC-PreInd-BF1_megahit_k127_179984_flag_0_multi_15.9657_len_40305</td><td>MC-PreInd-BF1_megahit_k127_179984_flag_0_multi_15.9657_len_40305_1</td></tr>
<tr><td>prophage.10</td><td>MC-PreInd-BF1_megahit_k127_233705_flag_0_multi_79.3140_len_11257|provirus_2183_11257</td><td>MC-12PostInd-BF1_megahit_k127_558051_flag_0_multi_34.4574_len_15183</td></tr>
<tr><td>prophage.11</td><td>MC-PreInd-BF1_megahit_k127_712980_flag_1_multi_9.0000_len_46935|provirus_2_40303</td><td>MC-PreInd-BF1_megahit_k127_712980_flag_1_multi_9.0000_len_46935|provirus_2_40303</td></tr>
</table>

**Table S4: IPHoP host matching information** | IPHoP host prediction for viruses of interest. The table includes the Viral genome ID, amino acid identity (AAI) to the closest RaFaH reference, predicted host genus, confidence score, and the list of methods used to assign host taxonomy.

| Viral genome ID | AAI to closest RaFAH ref | Host genus | Confidence score | List of methods |
|---|---|---|---|---|
| CT-12PostInd-BF1_megahit_k127_412173_flag_1_multi_6.0000_len_18991_1 | NA | Rickettsia | 90.3 | blast;92.80 |
| CT-PreInd-BF1_megahit_k127_349326_flag_1_multi_11.9815_len_62618\|provirus_2_29344 | 4.79 | Vibrio | 93.2 | blast;95.20 iPHoP-RF;80.60 |
| MC-12PostInd-BF1_megahit_k127_567814_flag_0_multi_29.0000_len_13848_extended_partial\|provirus_3_16519 | 25.62 | Vibrio | 92.6 | iPHoP-RF;94.70 |
| MC-PreInd-BF1_megahit_k127_179984_flag_0_multi_15.9657_len_40305_1 | 38.43 | Vibrio | 99.3 | blast;96.80 iPHoP-RF;95.10 RaFAH;90.10 CRISPR;88.00 |
| MC-PreInd-BF1_megahit_k127_233705_flag_0_multi_79.3140_len_11257\|provirus_2183_11257 | 75.19 | Vibrio | 92.1 | blast;96.80 iPHoP-RF;95.10 RaFAH;90.10 CRISPR;88.00 |
| MC-12PostInd-BF4_megahit_k127_317508_flag_0_multi_92.9303_len_13319_extended_partial | 14.68 | Aliivibrio | 90.3 | iPHoP-RF;92.80 |
| MC-12PostInd-BF3_megahit_k127_137292_flag_0_multi_26.9193_len_3350_extended_partial | 21.18 | Vibrio | 90.3 | blast;92.80 iPHoP-RF;85.40 |
| MC-12PostInd-BF1_megahit_k127_438300_flag_0_multi_15.9845_len_9826_extended_circular | 38.34 | Vibrio | 95.9 | blast;92.80 iPHoP-RF;85.40 |
| CT-12PostInd-BF1_megahit_k127_179695_flag_0_multi_155.5616_len_8741_extended_partial | 18.75 | Vibrio | 92.6 | iPHoP-RF;94.70 |
| MC-12PostInd-BF5_megahit_k127_220143_flag_0_multi_7.8228_len_3186_extended_partial | 21.13 | Cobetia | 90.1 | blast;92.60 iPHoP-RF;64.30 |
| MC-12PostInd-BF1_megahit_k127_47155_flag_0_multi_9.9100_len_10806 | 61.96 | Vibrio | 95.3 | iPHoP-RF;96.70 blast;93.10 |
| MC-12PostInd-BF1_megahit_k127_722844_flag_0_multi_11.0000_len_15850_extended_partial | 20.93 | Vibrio | 95.9 | iPHoP-RF;97.10 blast;92.30 |
| MC-12PostInd-BF1_megahit_k127_414575_flag_1_multi_8.8469_len_25558 | 46.86 | Vibrio | 95.4 | blast;96.80 iPHoP-RF;95.70 |
| CT-12PostInd-BF1_megahit_k127_414754_flag_0_multi_63.9153_len_33511_extended_partial | 16.05 | Vibrio | 92.2 | iPHoP-RF;94.40 |
| MC-12PostInd-BF1_megahit_k127_288805_flag_0_multi_26.0000_len_6225 | 97.4 | Vibrio | 100.0 | RaFAH;99.50 iPHoP-RF;96.70 blast;95.60 CRISPR;52.00 |

**Table S5: AphA BLASTp for proteins identified in the biofilm induction experiment (Prophage.8 and CT-PostInd-bin.5) and one virus identified in in situ *S. natans* samples (sarg_contig.k127_668789) |** BLASTp results for AphA proteins identified in this study against one another.

| Sequence ID | Query ID | % ID | Query coverage % | Query length | Sequence coverage % | Sequence length | Match Length | Mismatch | e-value |
|---|---|---|---|---|---|---|---|---|---|
| Prophage.8_17 | CT-PostInd-bin.5: k127_35206_8 | 34.78 | 90 | 187 | 84 | 160 | 161 | 84 | 5.68e-19 |
| sarg_contig.k127_668789_9 | CT-PostInd-bin.5: k127_35206_8 | 34.46 | 78 | 187 | 91 | 206 | 177 | 93 | 1.32e-19 |
| Prophage.8_17 | sarg_contig.k127_66879_9 | 99.38 | 100 | 206 | 78 | 160 | 160 | 1 | 6.14e-121 |
